# Neural mechanisms of the temporal response of cortical neurons to intracortical microstimulation

**DOI:** 10.1101/2023.10.15.562403

**Authors:** Karthik Kumaravelu, Warren M. Grill

## Abstract

**Background:** Intracortical microstimulation (ICMS) is used to map neuronal circuitry in the brain and restore lost sensory function, including vision, hearing, and somatosensation. The temporal response of cortical neurons to single pulse ICMS is remarkably stereotyped and comprises short latency excitation followed by prolonged inhibition and, in some cases, rebound excitation. However, the neural origin of the different response components to ICMS are poorly understood, and the interactions between the three response components during trains of ICMS pulses remains unclear.

**Objective:** We used computational modeling to determine the mechanisms contributing to the temporal response to ICMS in model cortical pyramidal neurons.

**Methods:** We built a biophysically based computational model of a cortical column comprising neurons with realistic morphology and synapses and quantified the temporal response of cortical neurons to different ICMS protocols. We characterized the temporal responses to single pulse ICMS across stimulation intensities and inhibitory (GABA-B/GABA-A) synaptic strengths. To probe interactions between response components, we quantified the response to paired pulse ICMS at different inter-pulse intervals and the response to short trains at different stimulation frequencies. Finally, we evaluated the performance of biomimetic ICMS trains in evoking a sustained neural response.

**Results:** Single pulse ICMS evoked short latency excitation followed by a period of inhibition, but model neurons did not exhibit post-inhibitory rebound excitation. The strength of short latency excitation increased and the duration of inhibition increased with increased stimulation amplitude. Prolonged inhibition resulted from both after-hyperpolarization currents and GABA-B synaptic transmission. During the paired pulse protocol, the strength of short latency excitation evoked by a test pulse decreased marginally compared to those evoked by a single pulse for interpulse intervals (IPI) <100 ms. Further, the duration of inhibition evoked by the test pulse was prolonged compared to single pulse for IPIs < 40ms and was not predicted by linear superposition of individual inhibitory responses. For IPIs>40 ms, the duration of inhibition evoked by the test pulse was comparable to those evoked by a single pulse. Short ICMS trains evoked repetitive excitatory responses against a background of inhibition. However, the strength of the repetitive excitatory response declined during ICMS at higher frequencies. Further, the duration of inhibition at the cessation of ICMS at higher frequencies was prolonged compared to the duration following a single pulse. Biomimetic pulse trains evoked comparable neural response between the onset and offset phases despite the presence of stimulation induced inhibition.

**Conclusions:** The cortical column model replicated the short latency excitation and long-lasting inhibitory components of the stereotyped neural response documented in experimental ICMS studies. Both cellular and synaptic mechanisms influenced the response components generated by ICMS. The non-linear interactions between response components resulted in dynamic ICMS-evoked neural activity and may play an important role in mediating the ICMS-induced precepts.

**HIGHLIGHTS:**

- Implemented a biophysically based computational model of the cortical column to study the temporal response of neurons to intracortical microstimulation (ICMS)
- Temporal response of model neurons comprised short latency excitation followed by a long-lasting inhibition but did not include rebound excitation.
- Excitation was mediated by both direct (antidromic) and indirect synaptic mechanisms and inhibition by both cellular (after-hyperpolarizing currents) and synaptic (GABAergic) mechanisms.
- The temporal dynamics of the response to ICMS should be considered when designing paradigms for sensory prosthetic applications.

## INTRODUCTION

Intracortical microstimulation (ICMS) is a powerful tool both to probe neural circuits [1–3] and to produce artificial sensations of a variety of modalities, including vision, audition, and somatosensation [4–9]. For example, ICMS has been explored to restore movement-related sensations such as touch and proprioception in persons with spinal cord injury [4, 7, 10]. The temporal response of cortical neurons in rodents and non-human primates (NHP) is remarkably stereotyped [1, 11, 12], and single pulse ICMS evokes short latency excitation (0-25 ms) followed by a period of inhibition (25-200 ms) and post-inhibitory rebound excitation (200-300 ms). However, the neural origins of this apparently ubiquitous response to single pulse ICMS are unclear and interactions with subsequent stimulus pulses during trains of stimulation remain poorly understood.

Butovas and colleagues quantified the temporal response to ICMS of neurons in the somatosensory cortex (S1) of rats [1, 13]. At stimulation threshold, single pulse ICMS evoked a short latency excitatory response followed by long-lasting inhibition [1], and a number of other groups subsequently replicated these findings [2, 11, 12, 14-22]. The strength of short-latency excitation increased, and the duration of the inhibition became longer, with increased stimulation intensity [11, 12]. Higher stimulation intensities or frequencies also precipitated rebound excitation following the inhibitory phase [1, 12]. Subsequent pharmacological experiments revealed that the ICMS-induced inhibitory responses were dependent on GABA-B receptors [23]. Despite this understanding, no computational model exists that reproduces the stereotyped temporal effects of ICMS. Further, current experimental techniques used to quantify the neural responses to ICMS, such as 2-photon calcium imaging and microelectrode recordings, have limitations including limited temporal resolution, and short latency responses may be obscured by stimulation artifacts [1, 24-26]. The relative contribution of various biophysical mechanisms including after-hyperpolarization (AHP) currents, short-term synaptic depression, and GABAergic synaptic transmission to the ICMS induced inhibitory response are not known.

We used computational modeling to quantify and deconstruct the mechanisms underlying temporal responses to ICMS. We implemented a biophysically-based computational model of cortical neurons with realistic morphologies adapted from the Blue Brain library [27, 28]. The model comprised five different cell types arranged vertically in a column with five layers [29], and each cell type included excitatory and inhibitory synaptic inputs from other cortical neurons. The results provide a mechanistic basis for the temporal responses to single pulse, paired pulses, and trains of ICMS and reveal the basis for the efficacy of so-called “biomimetic” pulse trains in evoking more sustained neural responses.

## METHODS

### Computational model of the cortical column

The computational model used in this study was developed in earlier work to study the spatial effects of ICMS and is described in detail elsewhere [29]. We implemented a computational model of a population of 6410 biophysically-based Hodgkin-Huxley style multi-compartment cortical neurons based on the Blue Brain cell library [27, 28]. The model comprised three parts: (1) single-cell cortical neurons with realistic axon morphology, (2) synapses distributed on the dendritic tree of each neuron, and (3) the model neurons with synapses arranged in a cortical column with dimensions derived from histological sections from the rhesus macaque. The model neurons included five different cortical cell types: Layer 1 neurogliaform cell with dense axonal arbor (L1 NGC-DA), Layer 2/3 pyramidal cell (L2/3 PC), Layer 4 large basket cell (L4 LBC), Layer 5 thick tufted pyramidal cell (L5 TTPC) and Layer 6 tufted pyramidal cell (L6 TPC) (Fig 1A). We incorporated synapses from the Blue Brain project to study the synaptic effects of ICMS [27]. For each cell type, the position of the postsynaptic compartments (on the dendritic tree) where an inbound synaptic connection was received was mapped in the Blue Brain project [27]. Fig 1A shows the locations of excitatory and inhibitory postsynaptic connections received by L5 PC cells from five different cell types (L1 NGC-DA, L2/3 PC, L4 LBC, L5 PC, and L6 PC). Neurons within the network were not interconnected, but rather isolated from one another, and were subjected to ongoing (intrinsic) synaptic inputs, synaptic events that were generated by stimulation, and the direct effects of stimulation.

**Figure 1:**
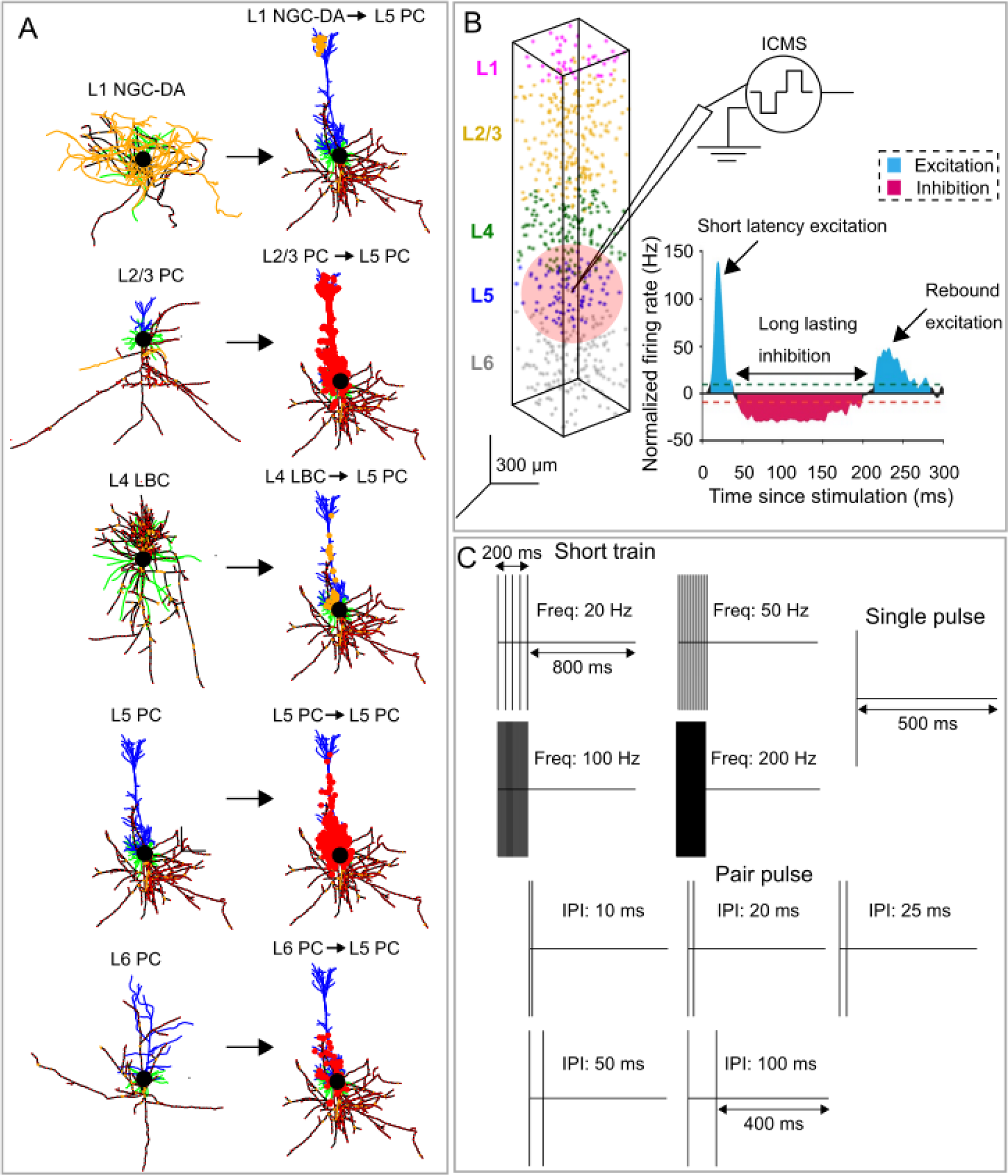
Biophysically-based computational model of intracortical microstimulation (ICMS) somatosensory cortex. (A) 3D morphology of Layer (L)-1 neuroglial cell, L2/3 pyramidal cell (PC), L4 basket cell, L5 thick tufted (TT) PC, L6 thick (T) PC. Model neurons were adapted from the Blue Brain library and had realistic axon morphologies [27, 28]. Blue – apical dendrites, green-basal dendrites, yellow-unmyelinated axon, black – myelin nodes, filled black circle-soma. Model axons were myelinated, and the diameter range was consistent with those found in rhesus macaques [28, 52]. Synaptic inputs to L5 PC from L1 NGC-DA, L2/3 PC, L4 LBC, L5 PC and L6 PC are shown in (A). Each cortical neuron (L1 NGC-DA, L2/3 PC, L4 LBC, L5 PC and L6 PC) received synaptic inputs from 55 presynaptic sources. All sources were cortical and did not include connections from other structures such as the thalamus, subcortex, etc. Excitatory and inhibitory postsynaptic compartments are shown in red and yellow-filled circles, respectively. Inhibitory synapses included GABA-A+GABA-B receptors and excitatory synapses included AMPA+NMDA receptors. The probability of synaptic activation to extracellular stimulation was an exponential function of the distance of the synapse from the electrode tip. (B) Cortical neurons were arranged in a column comprising five layers with dimensions of 400 µm x 400 µm x 2000 µm. Cortical thickness was determined from histological sections of the somatosensory cortex in the rhesus macaque [53], and the relative thickness of each layer was based on estimates from the visual cortex of rhesus macaque [54]. A uniform density of 20000 neurons/*mm*^3^ was used across all layers, resulting in a total cell count of 6410 neurons for the cortical column. The stimulation electrode was in L5, and the neural response was recorded from L5 PCs with somas located within 150 µm from the stimulation electrode. The light pink circle indicates the recording volume around the stimulation electrode. The inset shows the stereotyped temporal response to cortical stimulation recorded experimentally in L5 PC of rats including short latency excitation followed by a long-lasting inhibition and rebound excitation [16]. Dashed lines, significance thresholds determined by the 2.5 or 97.5 percentile of the empirical distribution of baseline normalized firing rate. (C) Three different ICMS protocols used in the model simulations: single pulse at different stimulation intensities, paired pulses at different inter-pulse intervals, and short trains at different stimulation frequencies.

Each postsynaptic compartment contained models of both AMPA and NMDA receptor kinetics for excitatory connections and GABA-A and GABA-B receptor kinetics for inhibitory connections, as described in the original publication [27]. In short, the synapses were modeled using a stochastic version of the Tsodyks-Markram model for dynamic synaptic transmission, including short-term facilitation (STF) and depression (STD). Fig 2 shows an example of synaptic dynamics (IPSC) for a GABA-B synapse in response to a 2, 20, 100 and 200 Hz stimulation train, and synaptic dynamics for the AMPA, NMDA and GABA-A receptors are shown in Fig S1-S3. The likelihood of activating a synapse by extracellular stimulation was an exponential function of the distance of the synapse from the electrode tip, and the space constant of the exponential function was larger for higher stimulation amplitudes than for lower amplitudes: 15 µA – 50 µm, 30 µA – 150 µm, 50 µA – 250 µm, 100 µA – 420 µm. Additional details on the derivation of spatial constants for synaptic activation are found elsewhere [29]. The cortical neurons with synapses were placed in a column with five layers (Fig 1B), and details of the cortical column, including dimensions, the relative thickness of each layer, neural density, etc., are provided elsewhere [29].

**Figure 2:**
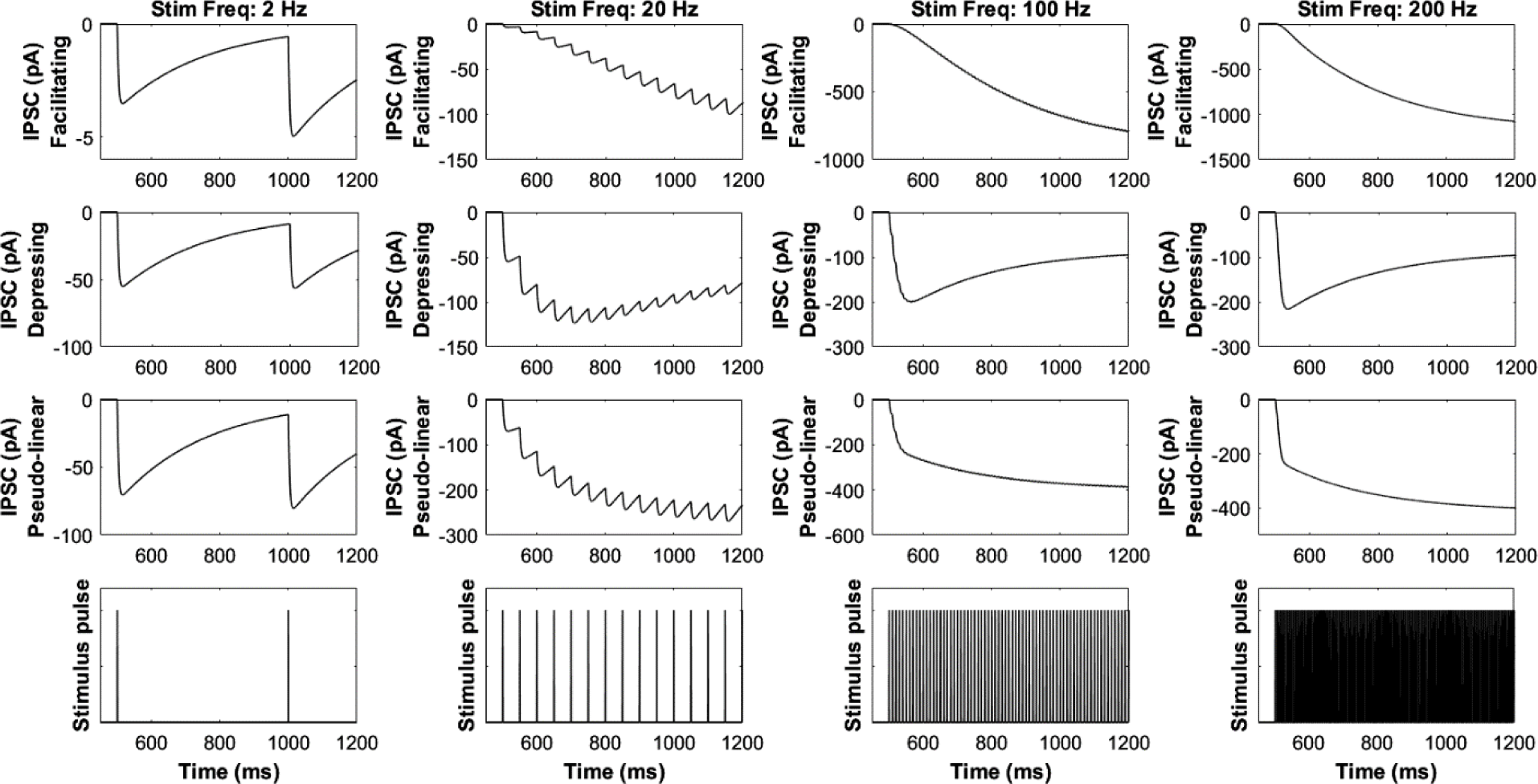
Properties of the (Tsodyks-Markram) GABA-B synapse. Inhibitory postsynaptic current (IPSC) showing dynamics of facilitating, depressing and pseudo-linear synapse in response to low (2, 20 Hz) and high (100, 200 Hz) frequency stimulation. Stimulation was applied from t=500ms. See Supplementary Figures S1-S3 for dynamics of other synapse types.

### Generation of intrinsic activity

To study the inhibitory effects of ICMS, the neurons must exhibit some ongoing intrinsic activity. We injected a 1.3 nA direct current (DC) intracellularly into the soma of each L5 PC, resulting in regular firing at a rate of ∼20 Hz. In a subset of simulations, we studied the effect of different baseline firing rates on ICMS-induced temporal responses by modulating the intensity of the injected DC (1-2 nA in steps of 0.5 nA). We also generated intrinsic activity by driving excitatory synapses on the L5 PC with 20 Hz Poisson spike trains. This gave rise to an irregular firing activity in the model neurons at an average frequency of ∼20 Hz. We also assessed the effect of the activation of different proportions of L5 excitatory synapses on the mean firing rate (Fig S4). Driving only 10% of excitatory synapses with 20 Hz Poisson spikes did not generate any intrinsic activity in the model neurons (Fig S4A). Driving 70%-100% of excitatory synapses with 20 Hz Poisson spikes yielded similar mean firing rates (13-21 Hz), with a 100% activation yielding a slightly higher firing rate than 70% (Fig S4). Both intracellular current injection and ongoing synaptic inputs were used independently to generate intrinsic activity to assess the impact of the baseline firing on neural activity induced by single pulse ICMS. Since both methods produced identical results, for the paired pulse and biomimetic protocols, intrinsic activity was generated only using DC current injection. For the short-train stimulation protocol, intrinsic activity was generated only using ongoing synaptic inputs.

### Neural data processing

Spiking activity was recorded from only those model neurons that were within a 150 µm radius around the stimulating electrode (Fig. 1B). Since the stimulating electrode was in L5 of the cortical column, activity was primarily recorded from L5 PCs. We constructed poststimulus time histograms (PSTH) by binning spike times in 5 ms bins across ten stimulation trials. We averaged the PSTH across ten stimulus trials for each neuron and then averaged the response across the neurons. The strength of the excitatory response was quantified by measuring the peak magnitude in the averaged PSTH response. Further, to quantify the relative contribution of direct versus synaptic activation to the excitatory response, we ran simulations of the model with and without synapses. We consider the relative contribution of direct activation to be peak excitatory response without synapses divided by the peak excitatory response in the model with synapses. For quantifying relative contribution of synaptic activation, we calculated the difference in peak excitatory response between the models with and without synapses and then divided that by the peak excitatory response in the model with synapses. Next, for each neuron, we quantified the duration of the inhibitory response by measuring the period when the firing rate was below 0.75 x the intrinsic firing rate for at least two consecutive bins. To quantify the relative contribution of AHP currents versus GABAergic synaptic transmission to the inhibitory response, we ran simulations of the model with and without inhibitory synapses. We consider the relative contribution of AHP currents to be inhibitory response duration without inhibitory synapses divided by the duration of inhibition in the intact model. For quantifying relative contribution of GABA synapses, we calculated the difference in inhibition duration between the models with and without inhibitory synapses and then divided that by the duration of inhibition in the model with synapses. For the short train protocol, a line was fit to the peak magnitude of the short latency excitatory responses evoked by each pulse in the stimulus train and the slope was estimated (*peak excitatiory response* (*Hz*) = *m* ∗ *time to peak* (*ms*) + *c*). To assess the variation of temporal response with cortical depth, stimulation was delivered in L2/3 and L6 for a subset of simulations, and recordings were made from neurons within a 150 µm radius around the stimulating electrode.

### ICMS protocols

We tested three ICMS protocols: single pulse, paired pulse, and short trains (Fig 1C). For single pulse and paired pulse protocols, ICMS was delivered at a repetition frequency of 2 Hz. The total simulation time was 5 s and with a repetition frequency of 2 Hz, this resulted in a total of 10 ICMS trials. For a subset of simulations, single pulse was delivered at repetition rates of 0.1, 0.5, 1 and 2 Hz. The total simulation time for each repetition rate was adjusted to get a total of 10 ICMS trials. For the short train protocol, ICMS was delivered at a repetition frequency of 1 Hz. The total simulation time was 10 s and with a repetition frequency of 1 Hz, this resulted in a total of 10 ICMS trials.

All stimuli were biphasic pulses (cathodic first) with a fixed width of 200 µs/phase and an interphase interval of 50 µs. The single pulse was delivered at four clinically relevant intensities (15, 30, 50 and 100 µA) [4]. The depth-dependent excitation thresholds from the model were compared to published depth-dependent detection thresholds from NHP studies (Fig S5-S6). Single pulse responses were also quantified at different GABA-B/GABA-A 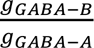 synaptic strength ratios (0.2-1 in steps of 0.2). For some simulations, responses were quantified with excitatory and inhibitory synapses turned off by setting the synaptic conductances to zero (i.e., g_syn=0 nS). For the paired pulse protocol, we quantified the response to five different interpulse intervals between the conditioning pulse and test pulse (10, 20, 25, 50 and 100 ms) at each stimulus intensity mentioned above. The timing of the test pulse always fell within the period of the prolonged inhibition. For the short train protocol, ICMS intensity was fixed at 50 µA, the frequency of pulses within the train was varied to include two low (20, 50 Hz) and two high frequencies (100, 200 Hz), and the duration of the short stimulus train was 200 ms. The duration of the short stimulus train was set to match those typically used in preclinical and clinical studies to induce sensory percepts [4, 7, 30].

### Biomimetic ICMS trains

Biomimetic ICMS trains are intended to evoke neural activity in somatosensory cortex that mimics that evoked by sensory inputs such as tactile stimuli [25, 31]. We tested biomimetic ICMS trains intended to mimic neural activity evoked by mechanical indentations applied onto the fingerpads of digits [32]. The mechanical stimulus consisted of 1 s long trapezoidal indentations delivered at a rate of 10 mm/s and depths ranging from 25-2000 µm (Fig. 10A). The somatosensory cortex is known to encode contact transients rather than the static contact [32], and thus ICMS trains were linearly mapped from the derivative of the indentation. The biomimetic ICMS trains consisted of five patterns with onset/offset phase durations of 10, 20, 50, 100 and 200 ms (Fig. 10A). The onset/offset phases comprised cathodic first biphasic pulses with a fixed width of 200 µs/phase, interphase interval of 50 µs, frequency of 300 Hz and an amplitude of 50 µA.

### Modeling ICMS

The stimulating tip of the electrode was approximated as a point current source [31, 33]. Extracellular potentials *V*_*e*_(*i*, *j*) due to the point current source in each compartment (*j*) of each model cortical neuron (*i*) were computed using the equation,

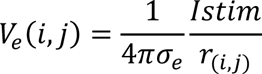

where *Istim* is the stimulating current, *r*_*i*,*j*_ is the distance from the stimulating electrode tip to each compartment of the model cortical neuron, and *σ*_*e*_ is the isotropic and homogeneous conductivity of the extracellular medium (0.3 S/m) [34]. All neurons that had a neural element (dendritic, somatic or axonal compartments) within 15 µm of the electrode tip were excluded from the analyses since *V*_*e*_ estimation using a point source would be inaccurate for such close distances without considering the 3D geometry of the microelectrode [33].

Simulations were implemented in NEURON 7.7 with equations solved using the backward Euler method with a time step of 0.025 ms [35]. The values of *V_e_* were coupled to each neuronal compartment using the *e_extracellular* mechanism [35]. The simulation was parallelized by distributing the total number of neurons (6410) across 50 processors in a round-robin fashion (128 neurons/processor) [36]. The code for the model required to replicate the results will be available on ModelDB post-publication.

### Statistical Analysis

Statistical tests were used to infer the effects of ICMS intensity on the duration of the inhibitory response. First, we tested the assumption of normality of inhibition duration at each ICMS intensity using the one-sample Kolmogorov-Smirnov test. Inhibition duration data were not normally distributed. Therefore, statistical inferences about the effect of ICMS intensity on inhibition duration were made using the Kruskal-Wallis one-way analysis of variance (ANOVA). When the omnibus test statistic revealed significance at *p<0.05*, we performed Dunn-Sidak’s test for *post hoc* paired comparisons between individual ICMS intensities [37]. Similar tests were used to infer the effects of ICMS frequency on the slope of the excitatory response. Further, one sample Wilcoxon signed-rank test was used to determine if the median slope was significantly different from zero. All statistical analyses were performed using MATLAB (Mathworks, Natick, MA).

## RESULTS

### Temporal response to single pulse ICMS

We quantified the response of model L5 cortical neurons to single pulse ICMS across different ICMS intensities and ratios of inhibitory synaptic conductances, 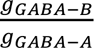. The response to single pulse ICMS included short latency excitation followed by a period of inhibition (Fig 3). ICMS in L2/3 or L6 generated similar excitation-inhibition responses in model L2/3 and L6 pyramidal neurons, respectively (Figs S7-S8). Both responses were observed for all stimulus strengths, and the strength of the short latency excitatory response and the duration of the inhibitory response increased with the intensity of stimulation (Fig 3C, D, E, F). At a 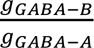 ratio of 0.6, the median inhibition duration across neurons in response to 15 µA ICMS was ∼10 ms and increased to ∼200 ms at 100 µA (Fig 4 B2). Stimulus intensity exerted a significant effect on inhibition duration (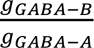 1, *p* = 10^−27^, *Kruskal – Wallis ANOVA*, χ^2^(3) = 126.65), and post hoc analysis revealed a significant difference in inhibition duration between all stimulation intensities (Fig 4, *p* < 0.05, *Dunn* – *Sidak method*). The strength of the short latency excitatory response did not change substantially with different 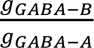 ratios (Fig 4 A1, B1, C1), but 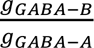 the duration of the inhibitory component increased with 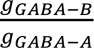 ratio (Fig 4). For instance, for a stimulus intensity of 30 µA, the median inhibition duration at 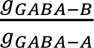 ratio of 0.2 was ∼85 ms compared to the inhibition duration of ∼125 ms at 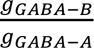 ratio of 1 (Fig 4). The temporal responses evoked by ICMS were similar whether the intrinsic activity was generated by DC current injection into the soma or via synaptic inputs (Figs 3, 4, S9, S10). The model neurons did not exhibit the post-inhibitory rebound excitation seen in experimental responses to ICMS at high intensities/frequencies (Figs 3, 4; see Discussion).

**Figure 3:**
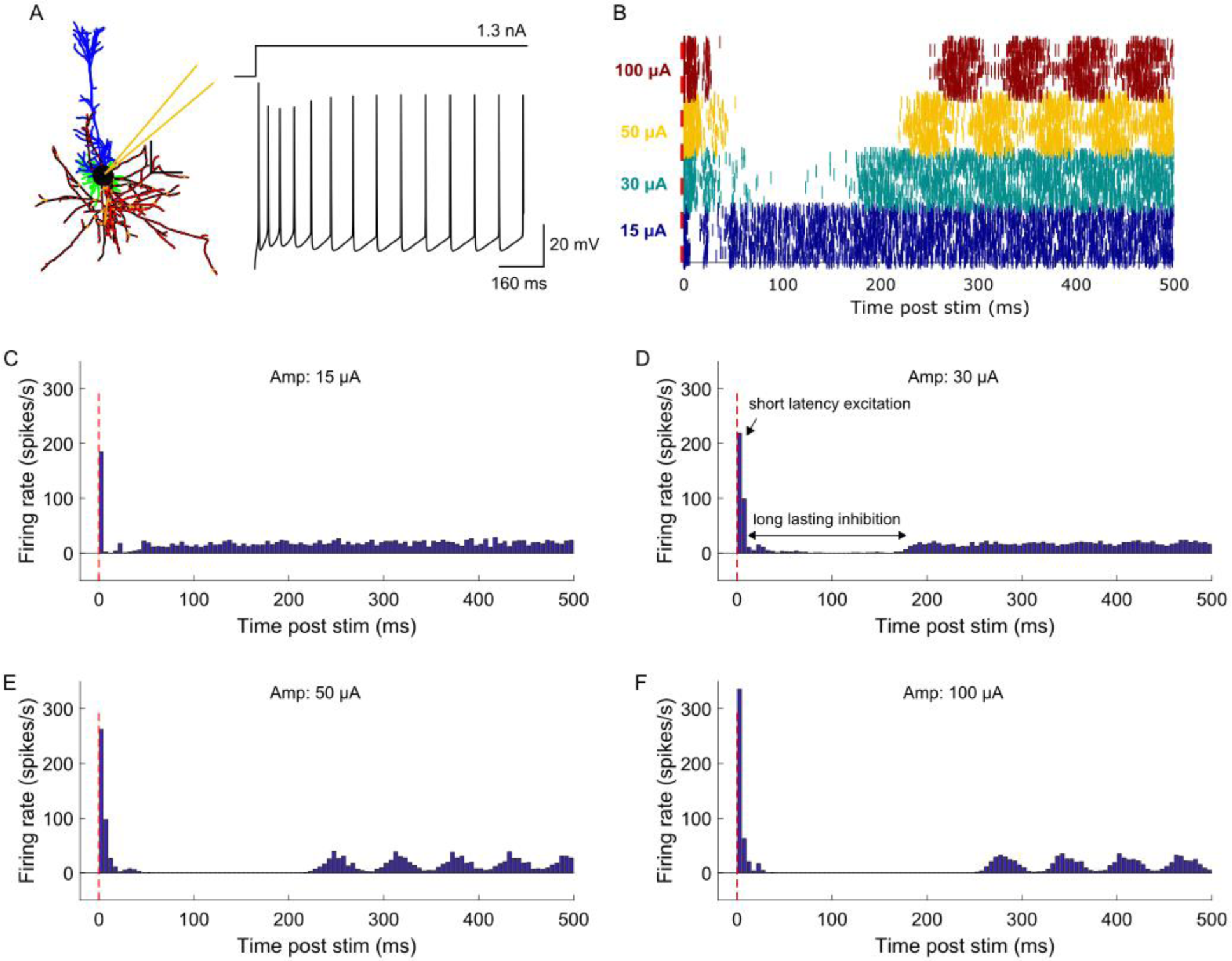
Temporal response of model neurons to ICMS at different stimulation intensities. (A) The intrinsic activity was generated by injecting a 1.3 nA DC current intracellularly into the soma of each neuron. (B) Raster plot showing firing activity of each of the individual 34 L5 PCs across stimulation intensities. Poststimulus time histogram (PSTH) response to ICMS at (C) 15 µA, (D) 30 µA, (E) 50 µA and (F) 100 µA. Stimulation delivered at 2 Hz comprised a single biphasic pulse (cathodic first) with a fixed width of 200 µs/phase and an interphase interval of 50 µs. The stimulation electrode was in L5. The response was averaged across 34 L5 PCs with somas located within 150 µm from the stimulation electrode. The 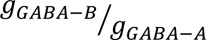 ratio was set at 1. A bin width of 5 ms was used to bin spike times. The temporal response to ICMS comprised short latency excitation followed by long-lasting inhibition. The strength of the excitatory response increased with stimulation intensity. Further, the duration of the inhibitory became longer with increased stimulation magnitude.

**Figure 4:**
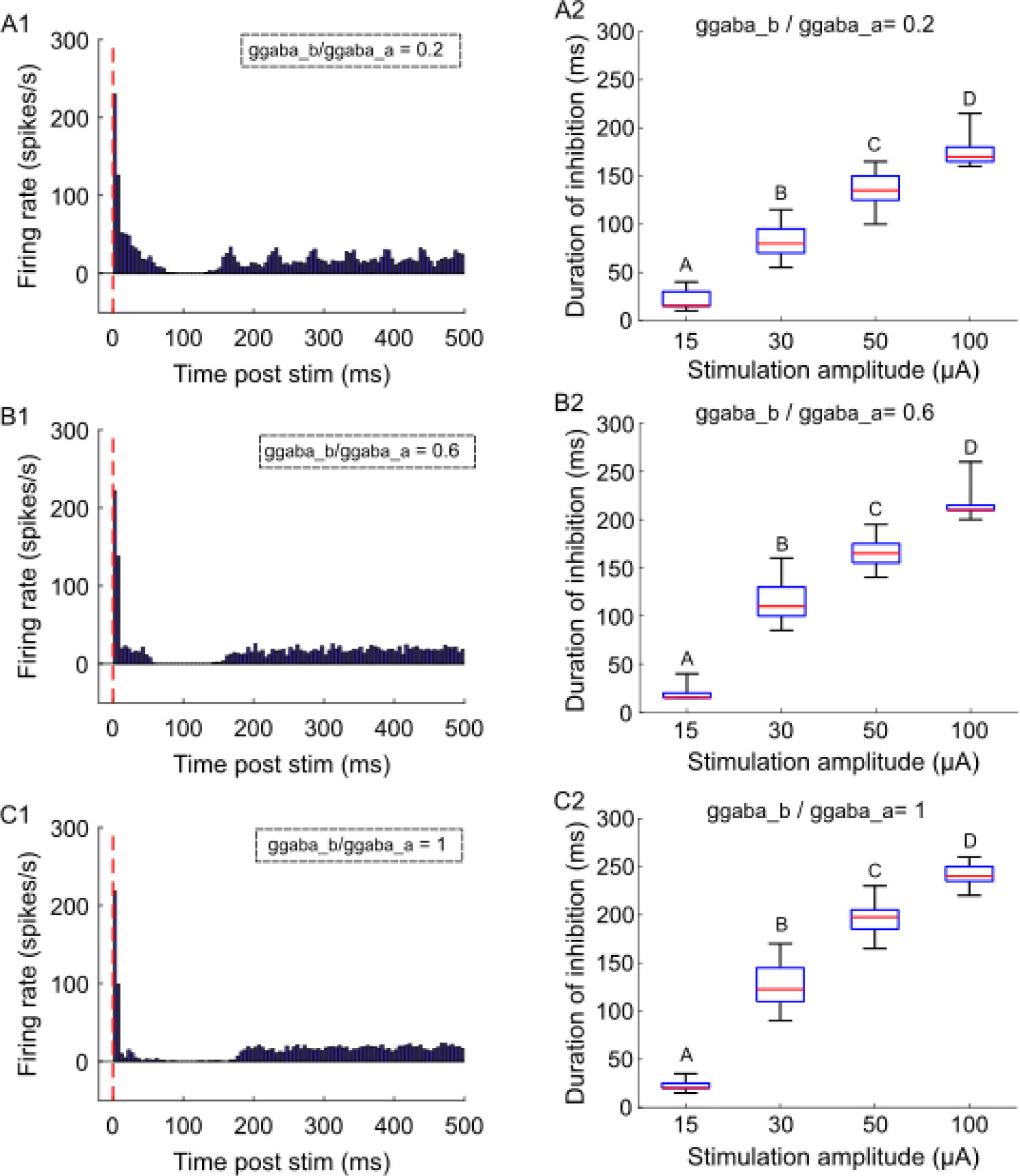
Model neuron responses to ICMS for different levels of GABA-B synaptic strength. PSTH response to ICMS at 30 µA and 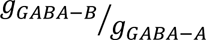 ratio of (A1) 0.2, (B1) 0.6, and (C1) 1. The response was averaged across 34 L5 PCs with somas located within 150 µm from the stimulation electrode. The intrinsic activity was generated by injecting 1.3 nA into the soma of neurons. A bin width of 5 ms was used to bin spike times. Duration of long-lasting inhibitory response as a function of ICMS amplitude for 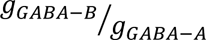 ratio of (A2) 0.2, (B2) 0.6, (C2) 1. Stimulation intensities that do not share the same letter are significantly different (p<0.05, Dunn-Sidak method). For each box, the central mark indicates the median duration across neurons, the bottom and top edges of the box indicate the 25th and 75th percentiles, and whiskers extend to 1.5 times the interquartile range. The duration of inhibition was shorter for weaker GABA-B synapses compared to stronger synapses. Further, the duration of inhibition increased with stimulation intensity.

To investigate the effects of the dynamic response evoked by one pulse on the responses evoked by subsequent pulses, we quantified the ICMS-induced neural activity with a single pulse delivered at different repetition rates – 0.1, 0.5, 1 and 2 Hz (Fig 5A, B, C, D). The strength of the excitatory response decreased marginally at a repetition rate of 2 Hz compared to the other rates of 0.1, 0.5 and 1 Hz (Fig 5A, B, C, D, E). The duration of the inhibitory response remained comparable across all repetition rates (Fig. 5F).

**Figure 5:**
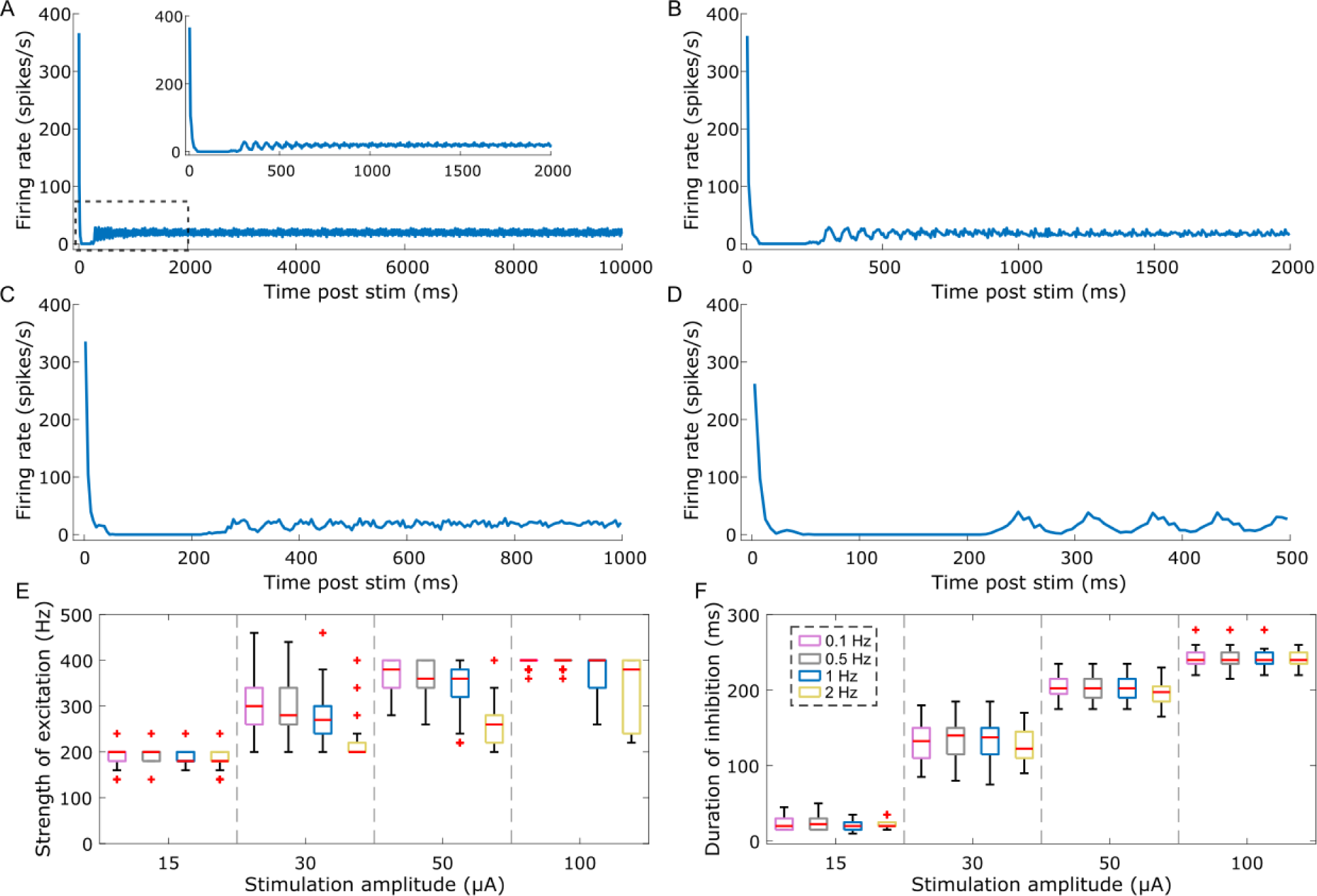
Model neuron response to single pulse ICMS delivered at different repetition rates. Stimulus triggered response to 50 µA ICMS pulse delivered at (A) 0.1 Hz, (B) 0.5 Hz, (C) 1 Hz and (D) 2 Hz. The inset in panel A shows the response to 0.1 Hz stimulation at a shorter time scale. For each condition, the stimulus triggered response was averaged across 10 pulses. The 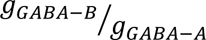 ratio was set at 1. The model neurons exhibited oscillatory activity post ICMS induced inhibitory response. Refer to Supplementary Figure S11 for response to 100 µA ICMS pulse at different repetition rates. (E) The peak of the excitatory response as a function of stimulation amplitude across the four repetition rates. (F) Duration of inhibition as a function of stimulation amplitude for the four repetition rates. Each color box represents a different repetition rate: 0.1 Hz (magenta), 0.5 Hz (gray), 1 Hz (blue), 2 Hz (yellow). For each box, the central mark indicates the median slope across neurons, the bottom and top edges of the box indicate the 25^th^ and 75th percentiles, whiskers extend to 1.5 times the interquartile range, and the plus signs indicate outliers.

After the stimulation induced excitatory-inhibitory response, the model neurons did not return to the ∼20 Hz regular firing baseline activity. Instead, there was an intensity-dependent modulation of the baseline activity – with response to low-intensity stimulation returning to baseline activity, whereas high-intensity stimulation generated a ∼20 Hz sinusoidal oscillatory activity (Fig 3C, D, E, F). This sinusoidal oscillation lasted for ∼450 ms before waning at the 50 µA stimulus intensity (Fig 5A, B, C, D). However, at the 100 µA intensity, the sinusoidal oscillatory activity did not cease to exist at even the lowest tested repetition frequency of 0.1 Hz (Fig S11). This modulation occurred only for regular firing baseline activity and not for irregular inputs (Figs 3C-F, S9C-F).

Next, we assessed whether the inhibitory response in the model neurons resulted from cellular mechanisms (afterhyperpolarization (AHP) currents) and/or synaptic mechanisms (GABA synapses and/or neurotransmitter depletion due to short-term depression). The inhibition duration across repetition rates remained consistent, indicating a change in excitatory synaptic conductance over time (i.e., short-term depression) not to affect the inhibitory response (Fig 5F). Removing inhibitory synaptic input increased the excitability of the model neurons, thereby increasing the strength of the short latency excitatory response compared to the control (with inhibitory synapses) condition (Fig 6A, B, E). Despite the lack of inhibitory synapses, the excitatory response was followed by a strong inhibitory response due to the AHP currents (Fig 6A, B). However, the duration of the inhibitory component condition was shorter with inhibitory synapses off than in the control condition (Fig 6D). Further, when both excitatory and inhibitory synapses were turned off, the strength of the excitatory response and the duration of the inhibitory response were both substantially reduced compared to the control condition (Fig 6D, E). These results were consistent across 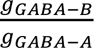 ratios (Fig S12-S13). For stimulation intensities of 15, 50 and 100 μA, AHP currents contributed strongly (>70%) to the inhibitory response compared to GABA synapses (<30%) (Fig 6F). However, for the stimulation intensity of 30 μA, both AHP currents and GABA synapses contributed equally to the stimulation-induced inhibitory response (Fig 6F). The excitatory response resulted mainly due to direct activation for stimulation intensities up to 50 μA, while both direct and synaptic activation contributed equally at 100 μA (Fig 6G).

**Figure 6:**
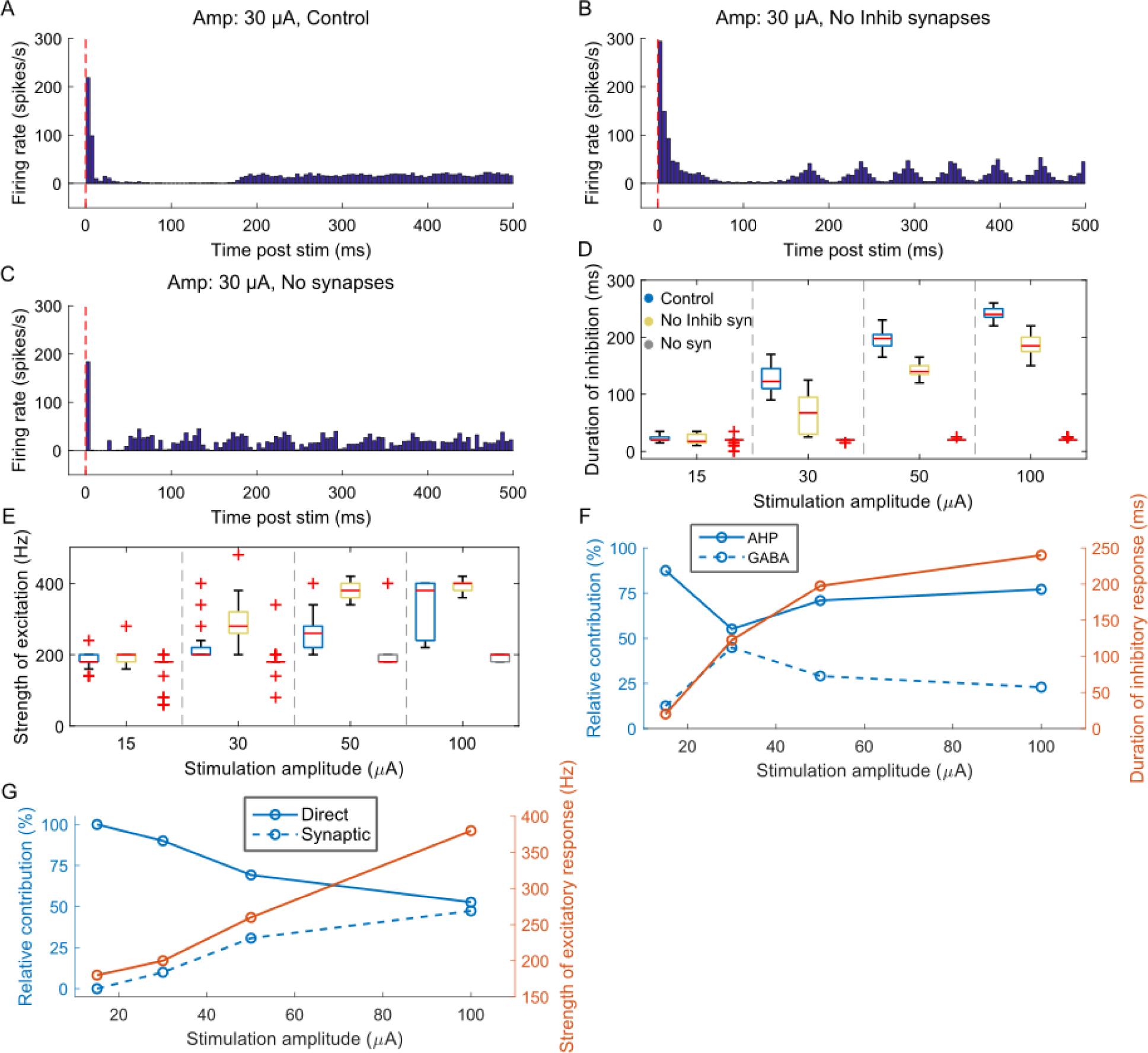
Temporal response of model neurons to ICMS at 30 µA and 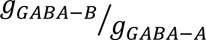 ratio of 1 for (A) control condition (intact synapses), (B) without inhibitory synapses, (C) without inhibitory and excitatory synapses. The response was averaged across 34 L5 PCs with somas located within 150 µm from the stimulation electrode. A bin width of 5 ms was used to bin spike times. (D) Duration of inhibition as a function of stimulation amplitude for the three conditions. (E) The peak of the excitatory response as a function of stimulation amplitude for the three conditions. Each color box represents a different simulation condition: control condition (blue), without inhibitory synapses (yellow), without inhibitory and excitatory synapses (gray). For each box, the central mark indicates the median slope across neurons, the bottom and top edges of the box indicate the 25th and 75th percentiles, whiskers extend to 1.5 times the interquartile range, and the plus signs indicate outliers. Refer to Supplementary Figures S12-S13 for other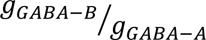 ratios. (F) Relative contribution of afterhyperpolarization (AHP) currents and GABA synapses to ICMS-evoked inhibitory response. (G) Relative contribution of direct vs. synaptic activation to ICMS-evoked short latency excitatory response.

Finally, we assessed the impact of different mean baseline firing rates on the responses evoked by single pulse ICMS. The strength of the excitatory response increased with baseline firing rates, and the duration of the inhibitory response decreased with an increase in the baseline firing (Fig 7). The mean baseline firing rate influenced the sinusoidal oscillatory activity that occurred after the stimulation induced excitatory-inhibitory response with lower rate (∼12 Hz) generating a sinusoidal response, while the higher rates did not (Fig 7).

**Figure 7:**
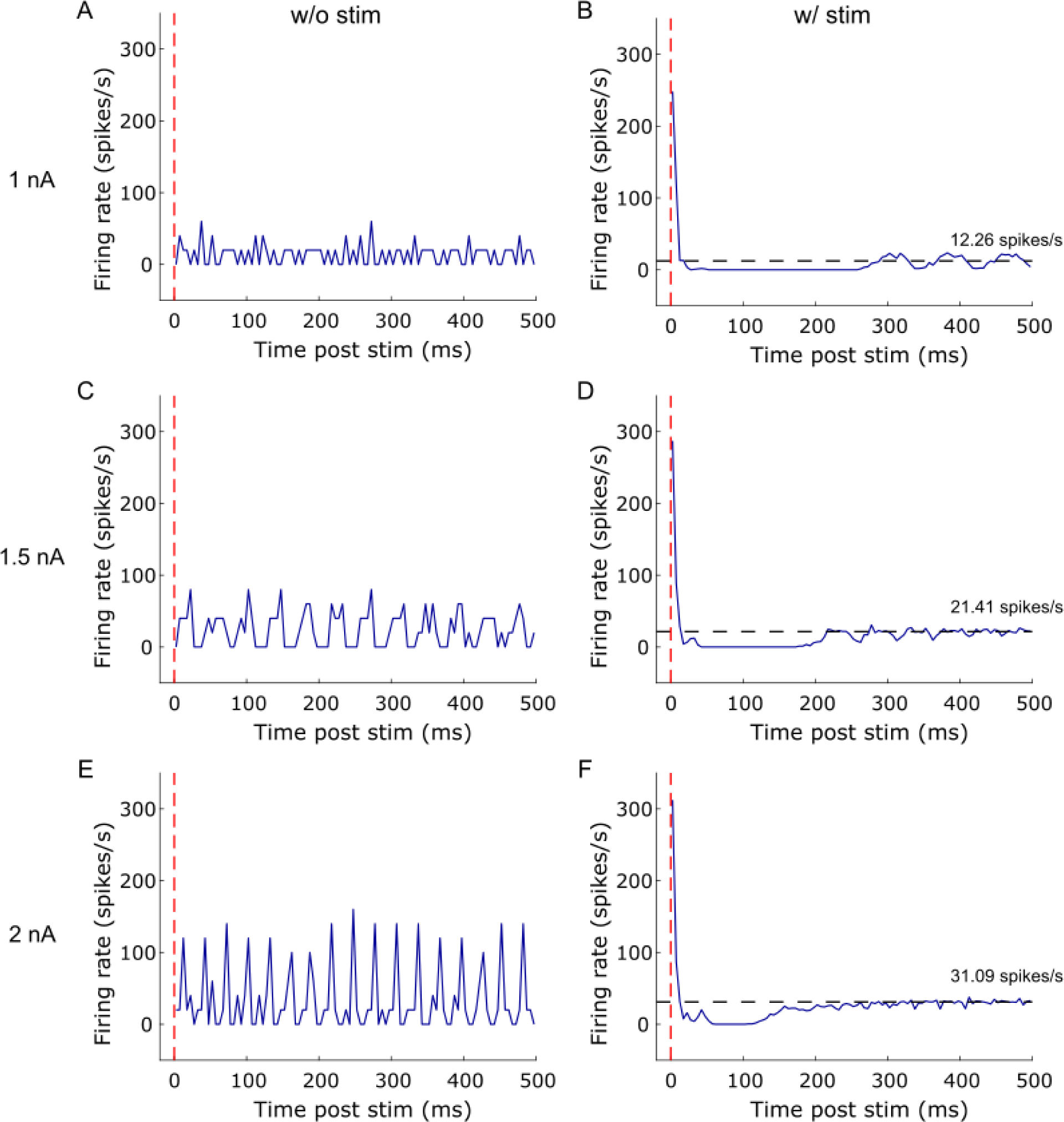
Effect of different mean intrinsic firing rates on the temporal response to ICMS at 50 μA and 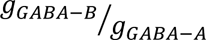 ratio of 1. Stimulus triggered response without ICMS for intracellular current injection of (A) 1nA, (C) 1.5 nA, (E) 2nA. Injecting a high DC into soma generated oscillatory baseline activity. Stimulus triggered response with ICMS for current injection levels of (B) 1nA, (D) 1.5nA, (F) 2nA. Increasing current injection levels yielded higher mean intrinsic firing rate indicated by the dashed black line. The duration of the inhibitory response is reduced with an increased mean firing rate. Further, the magnitude of the short latency excitatory response increased with the firing rate. At the lower baseline firing rate, the model neurons exhibited oscillatory activity after the inhibitory response.

### Temporal response to paired pulse ICMS

We delivered paired pulse ICMS at different interpulse intervals to investigate the effects of the inhibitory response evoked by the first (conditioning) pulse on the response evoked by the subsequent (test) pulse. The test pulse consistently evoked short latency excitatory and longer latency inhibitory responses across all IPIs (Fig 8A, B, C, D). The strength of the excitatory response decreased slightly, in comparison to the response evoked by a single pulse, independent of the IPI (Fig 8E). The duration of the inhibitory response evoked by the test pulse was comparable to that evoked by a single pulse for IPIs>40 ms but for IPI < 40 ms, the duration of inhibition evoked by the test pulse was longer than predicted by linear superposition of the duration evoked by a single pulse plus the IPI. (Fig 8F). These responses to the paired pulse protocol were consistent across stimulation intensities (Fig S14, S15).

**Figure 8:**
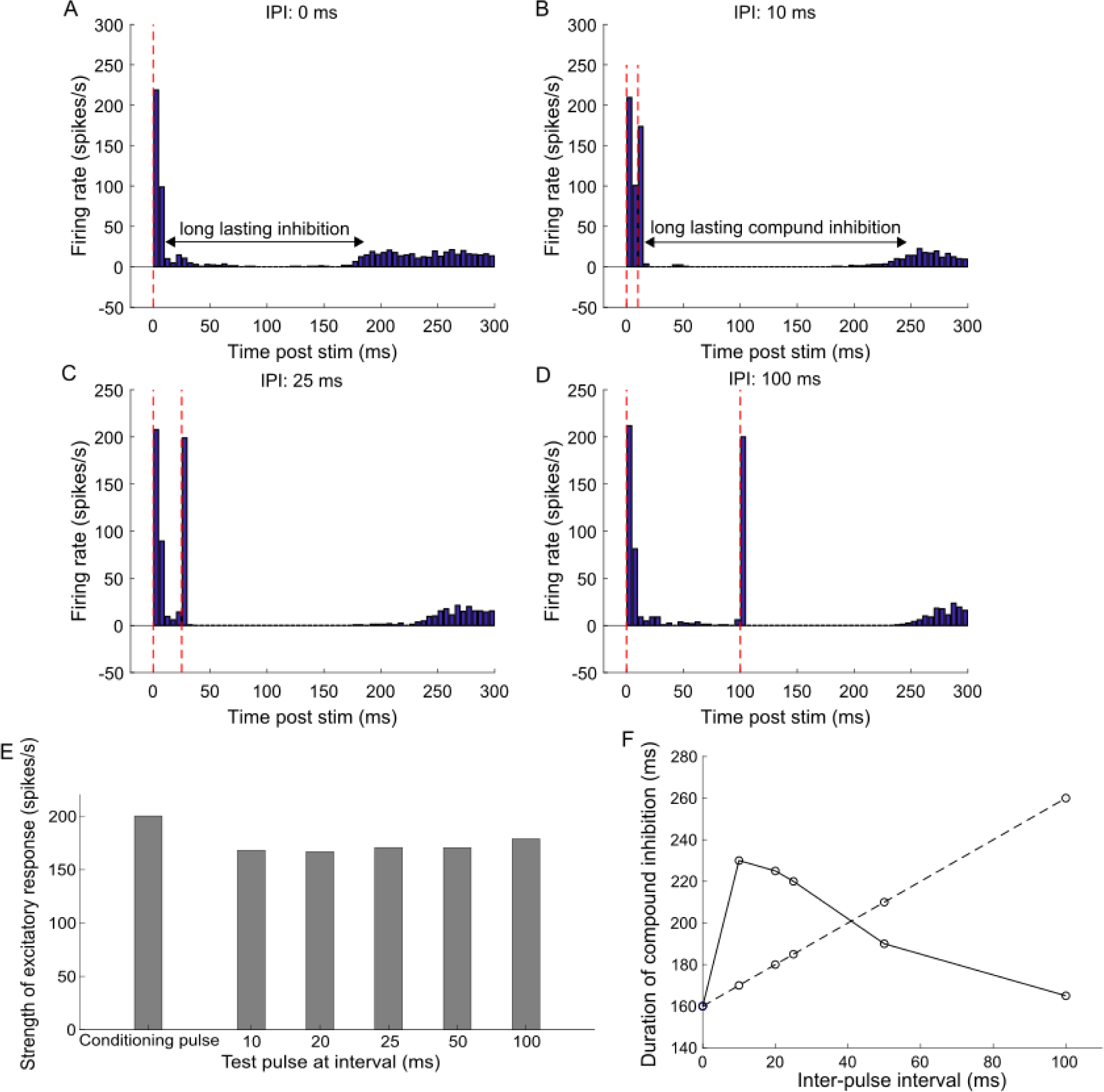
Response of model neurons to paired pulse ICMS. (A) PSTH response to a single (conditioning) pulse applied at *t* = 0 *ms*. Response to paired pulses with conditioning pulse applied at *t* = 0 *ms* and test pulse applied at an inter-pulse interval (IPI) of (B) 10 ms, (C) 25 ms, (D) 100 ms. The red dashed lines indicate the timing of the control and test pulses. The paired-pulse stimulation was applied at 2 Hz and 30 µA. The conditioning pulse was applied at the same amplitude as the test pulse. Further, the interval of the test pulse was chosen to fall within the duration of the inhibitory response generated by the control pulse. The 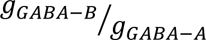 ratio was set at 1. A bin width of 5 ms was used to bin spike times. (E) Strength of short latency excitatory response to test pulse at 30 µA. The strength of excitatory response to test pulse decreased marginally compared to a single (conditioning) pulse and this effect was consistent across all stimulation intensities. This decrease can be attributed to the inhibitory effect triggered by the conditioning pulse. (F) The duration of inhibitory response to test pulse at 30 µA. Solid trace shows the inhibition duration to test pulses, whereas the dashed line indicates the duration predicted by linear superposition of individual inhibitory responses, i.e., duration of inhibition following single pulse + IPI. There was supra-linear addition of individual inhibitory responses for IPIs ≤ 40 ms and a sub-linear addition for IPIs >40 ms.

### Temporal responses to short trains of ICMS

We applied short trains of ICMS, and the responses of model neurons were classified into three types based on the dynamics of the excitatory response during the stimulus train: (1) facilitation – the magnitude of the excitatory response increased during the course of the train (Fig 9A), (2) no change in the magnitude of excitatory responses during the train (Fig. 9B), or (3) depression – the magnitude of the excitatory responses decreased during the train (Fig. 9C). A line was fit to the peak excitatory response evoked by each pulse, and the slope was computed. Most neurons exhibited a negative slope (depression) at higher stimulus frequencies in contrast to lower frequencies, and the median slope across 34 L5 PC neurons was ∼-0.4 Hz/ms at 200 Hz compared to ∼0.1 Hz/ms at 20 Hz (Figure 9G). Stimulus frequency exerted a significant effect on the slope (*p* = 10^−9^, *Kruskal* − *Wallis ANOVA*, χ^2^(3) = 42.43), and *post hoc* analysis revealed a significant difference in slope across stimulation frequencies (Fig 9G, *p* < 0.05, *Dunn* − *Sidak method*). As well, for each stimulus frequency, a one-sample Wilcoxon signed-rank test indicated that the median slope was significantly different from 0 (20 Hz: Z = 4.5649, p = 4.9981e-06, 50 Hz: Z = 4.5179, p = 6.2454e-06, 100 Hz: Z = 3.0110, p = 0.0026, 200 Hz: Z = −3.8567, p = 1.1494e-04). These results suggest that neurons were better able to follow low stimulation frequencies than higher frequencies. Excitatory synapses mediated the facilitatory effects at low stimulation frequencies, whereas GABA synapses were involved in the depression at higher stimulation frequencies (Fig 9G). Further, the duration of inhibition at the end of the stimulus train at higher frequencies was longer than the duration of inhibition evoked by a single pulse (Fig. 9D, E, F). GABA synapses contributed strongly to the inhibitory response at low stimulus frequencies, while at higher frequencies (>=100 Hz) both AHP currents and GABA synapses contributed equally (Fig. 9H). These effects were consistent across 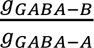 ratios (Fig S16).

**Figure 9:**
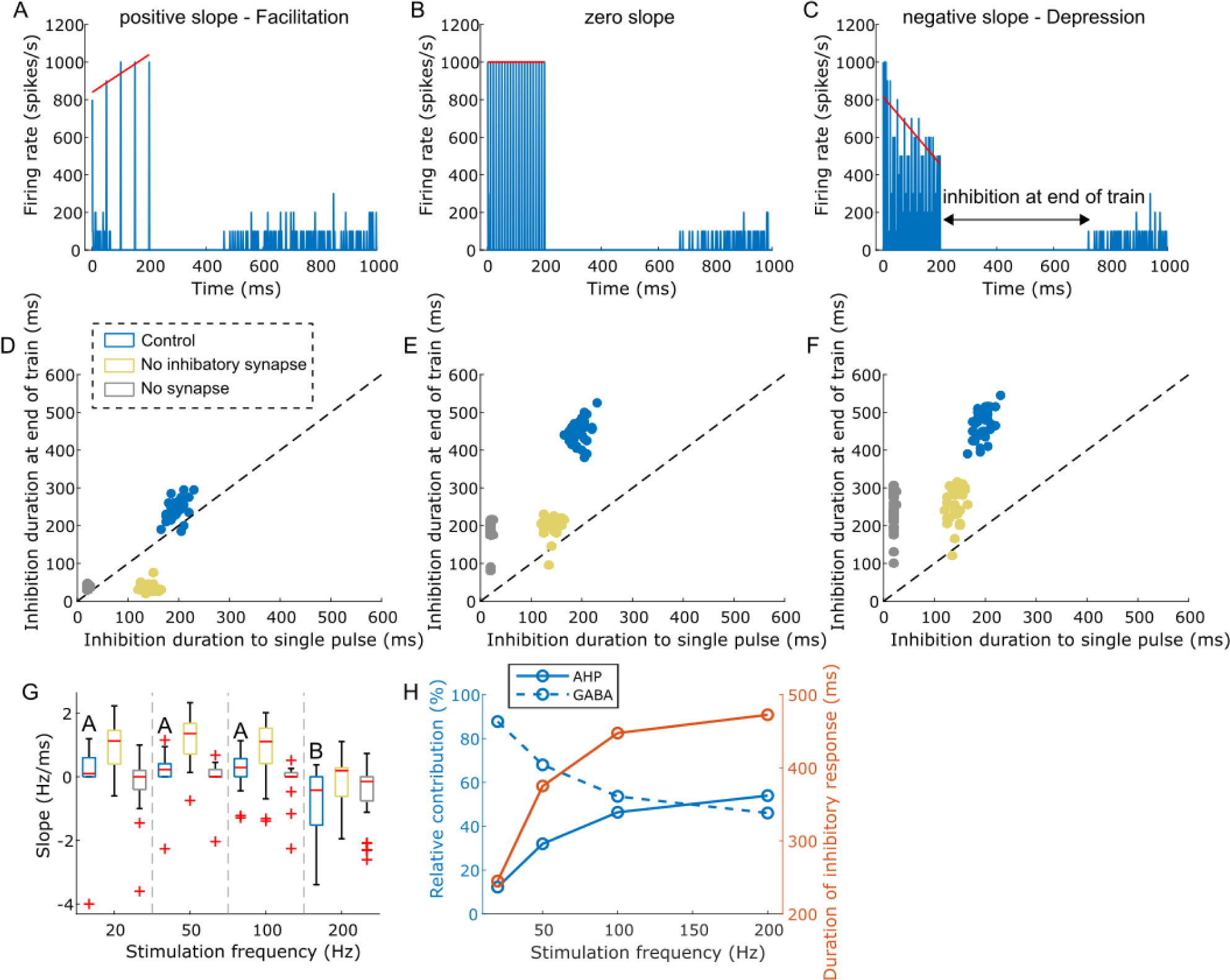
Model response to short trains of ICMS at different frequencies. A line was fit to the peak short latency excitatory responses evoked by each pulse within the stimulus train. Model neurons exhibited three types of response dynamics: (A) Positive slope - facilitation of the short latency excitatory response with the progression of the ICMS train, (B) Zero slope – no change in the excitatory response during the train, (C) Negative slope – decline of the short latency excitatory response with the progression of the ICMS train. A bin width of 1 ms was used to bin spike times for panels A-C. The 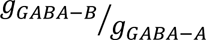 ratio was set at 1. Correlation of inhibition duration at the end of the train with inhibition duration following single pulse stimulation for short trains at frequencies (D) 20 Hz, (E) 100 Hz, (F) 200 Hz. Each dot represents the duration of inhibition of each 34 L5 PCs across three conditions: control condition (blue), without inhibitory synapses (yellow), without inhibitory and excitatory synapses (gray). The inhibition duration at the end of the ICMS train was longer than the duration following a single pulse for trains at higher stimulus frequencies compared to lower ones. (G) The slope of the line fit to the excitatory response as a function of stimulation frequency. For the control condition, stimulation frequencies that do not share the same letter are significantly different (p<0.05, Dunn-Sidak method). For each box, the central mark indicates the median slope across neurons, the bottom and top edges of the box indicate the 25th and 75th percentiles, whiskers extend to 1.5 times the interquartile range, and the plus signs indicate outliers. Each color box represents a different simulation condition: control condition (blue), without inhibitory synapses (yellow), and without inhibitory and excitatory synapses (gray). Most neurons exhibited a negative slope at higher stimulation frequencies. Refer to Supplementary Figure S16 for other 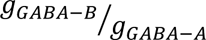 ratios. (H) Relative contribution of AHP currents and GABA synaptic transmission to ICMS-induced inhibitory response.

### Temporal responses to biomimetic ICMS trains

Patterns of stimulation in biomimetic pulse trains are intended to mimic the neural activity evoked during natural physiological inputs (Fig 10A). We quantified the temporal response to biomimetic ICMS trains to determine whether biomimetic stimuli can reliably evoke neural activity matching those evoked by mechanical (tactile) stimuli (Fig 10A). The biomimetic trains comprised an onset phase, a plateau phase, and an offset phase (Fig 10A). The biomimetic trains reliably evoked a series of short latency excitatory responses during the stimulus onset and offset (Fig. 10B1-B5), and the area under the curve of the response did not change substantially between the onset and offset phases (Fig 10C). The inhibition duration evoked at the end of the offset phase was similar to the duration at the end of the onset phase across all tested biomimetic stimulus trains (Fig. 10B1-B5).

**Figure 10:**
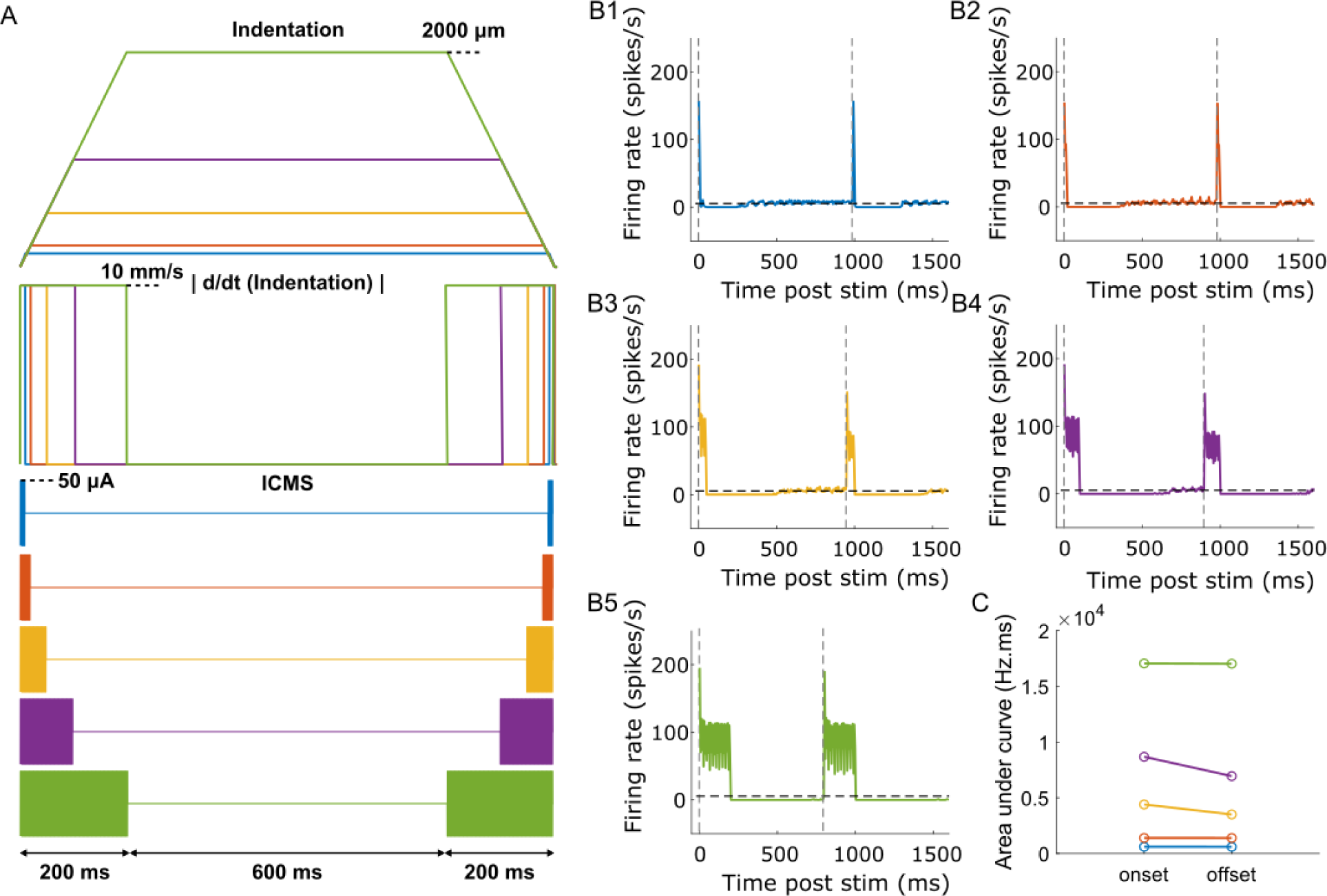
Temporal response of model neurons to biomimetic ICMS trains. (A) 1-s long trapezoidal indentations delivered at a rate of 10 mm/s and depths ranging from 25-2000 µm. Absolute value of the first derivative of trapezoidal indentations |*d*/*dt* (*indentation*)|. ICMS trains linearly mapped from |*d*/*dt* (*indentation*)|. The onset/offset phases comprised cathodic first biphasic pulses with a fixed width of 200 µs/phase, interphase interval of 50 µs, frequency of 300 Hz and an amplitude of 50 µA. The temporal response of model neurons to biomimetic ICMS trains with onset/offset phase duration of (B1) 10 ms, (B2) 20 ms, (B3) 50 ms, (B4) 100 ms, (B5) 200 ms. The 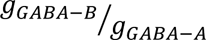 ratio was set at 1. The horizontal dashed black line indicates the mean intrinsic firing rate and vertical gray dashed line indicates onset of stimulation phases. A bin width of 5 ms was used to bin spike times. (C) Comparison of area under the curve of the responses during the onset and offset phases of the biomimetic ICMS trains.

## DISCUSSION

We implemented a computational model of a cortical column to deconstruct the mechanisms underlying the temporal responses to various ICMS protocols. Single pulse ICMS evoked a short latency excitatory response followed by a period of inhibition. Further, the strength of the excitatory response and the duration of inhibition increased with stimulation intensity. The excitatory response was primarily generated by direct activation at the lower stimulation intensities, while both direct and synaptic activation contributed equally at the higher stimulation intensities, and both AHP currents and GABAergic synaptic transmission were prominent in generating the inhibitory response.

GABAergic synaptic transmission generated a strong inhibitory response that lasted for several hundred milliseconds post-stimulation and was primarily responsible for the rapid decline of the short latency excitatory response during stimulus trains at higher frequencies. Finally, biomimetic stimulus trains evoked similar responses across the onset and offset phases despite the effect of GABAergic-induced inhibition.

### Model validation

The temporal response to ICMS has been recorded and characterized in the sensorimotor areas of both rodents and non-human primates [1, 11, 12, 14, 15, 18, 22]. Butovas and colleagues observed short latency excitation and long-lasting inhibition that were linked across stimulation intensities (0.8-4 nC) and rebound excitation occurred at higher stimulation intensities (>2 nC) [1]. Similarly, our model neurons exhibited short latency excitation and long-duration inhibition, but none of the stimulation intensities evoked rebound excitation. Several studies replicated the results of Butovas and Schwarz and observed an increase in the duration of inhibition with stimulus amplitude [11, 12, 20]. We found a significant increase in inhibition duration with the stimulation intensity. Butovas et al. conducted follow up studies using pharmacological blockers to probe the neural origin of the long-duration inhibitory response [23]. The inhibitory response to ICMS was reduced following block of GABA-B receptors, and our finding that the duration of the inhibitory response in model neurons increased with the GABA-B/GABA-A synaptic strength ratio is consistent with these results. Further, the state dependence of the ICMS evoked oscillatory response in the model agreed well with the results reported in a study by Venkatraman and Carmena [38]. In their study, ICMS during quiet trials (i.e., a low baseline firing rate) evoked an initial excitation response followed by prolonged inhibition and oscillatory bursting, similar to our model response [38]. However, ICMS delivered during trials with whisker deflection (i.e., high baseline firing rate) induced only an initial excitatory response followed by a short period of inhibition, consistent with our results [38]. Other cortical stimulation modalities such as transcranial magnetic stimulation (TMS) also evoke a triphasic temporal response pattern similar to ICMS [16, 17, 39], and the stimulus intensity dependent effects of TMS on the different response components are similar to the intensity dependent effects of ICMS [16, 17, 39].

In addition to the response to single pulse ICMS, Butovas and Schwarz quantified the response to paired pulses at IPIs of 25, 50 and 100 ms [1]. They found a supralinear addition of inhibition duration for the IPI of 25 ms and sublinear addition for the 50 and 100 ms IPIs. This matched with our findings where we observed a supra-linear addition for IPIs up to 40 ms and a sub-linear addition for IPIs >40 ms. The strength of the excitatory response to the test pulse at the various IPIs did not change substantially compared to the single pulse in the Butovas and Schwarz study, which was consistent with our findings.

Butovas and Schwarz also quantified the response to short stimulus trains at frequencies 5-40 Hz [1]. They found that low stimulation frequencies evoked repetitive excitatory responses over a background of inhibition, and we observed the same pattern in the model. In addition, we found that the excitatory response of most model neurons declined during the train at higher stimulus frequencies, similar to the results of recent studies [12, 40]. As well, Sombeck et al. observed that the duration of inhibition at the end of the train was longer than the inhibition following a single pulse for stimulus trains at higher frequencies than at lower frequencies, consistent with our results [12].

### Neural mechanisms of temporal response to single pulse ICMS

ICMS preferentially activates the axonal terminals near the stimulating microelectrode at the stimulus threshold [28, 29, 41]. The neural origin of the short-latency excitation response includes activation through antidromic (direct) and orthodromic (synaptic) propagation from activated axonal terminations. The excitatory response in the model neurons was predominantly generated by direct activation at lower stimulation intensities (15, 30 and 50 µA). However, both direct and synaptic activation contributed equally to the excitatory response at 100 µA. There are several potential contributors to the inhibitory response including AHP currents, GABAergic synaptic transmission, and short-term synaptic depression [42, 43]. We ruled out the possibility of short-term depression of excitatory neurotransmitters because the inhibitory response was comparable across repetition rates, indicating that the change in excitatory synaptic conductance over time had a limited effect on the inhibitory response. There was a non-monotonic relationship between the relative contributions of AHP currents and GABA synaptic transmission and the stimulus intensity. The AHP currents mainly contributed to the inhibitory response at stimulus intensities 15, 50 and 100 µA, while both AHP currents and GABA synaptic transmissions contributed equally to the inhibitory response at an intensity of 30 µA. The strength of the excitatory response (which generates the AHP currents) did not change much between stimulation intensities of 15 and 30 µA (refer Fig 6E). However, the space constant of GABA synaptic activation to ICMS tripled from 50 µm at 15 µA to 150 µm at 30 µA. This is why both AHP and GABA synaptic transmissions contributed equally to the inhibitory response at 30 µA. However, the strength of the excitatory response increased with stimulation intensity for 50 and 100 µA, thereby AHP currents dominating the generation of the inhibitory response at those intensities. Our findings on the neural origin of the temporal response to ICMS are expected to apply to the other cortical stimulation modalities, such as TMS, that generate a similar neural response to stimulation.

### Relationship between ICMS-induced neural activity and ICMS-induced sensory perception

It is not clear what role each component of the temporal response to ICMS contributes to induced behavioral effects or perception. The intensity of the tactile percepts induced by continuous ICMS declined more rapidly (tens of seconds) at higher stimulus frequencies than at lower ones [44]. Similarly, high frequency ICMS delivered at 100-200 Hz led the induced phosphenes to disappear within several hundred milliseconds [45]. The model suggests that this effect during continuous high frequency stimulation is due to the decline of the short latency excitatory response, which was more pronounced at higher frequencies. The literature includes several speculations as to why high frequency continuous ICMS may not be able to evoke a consistent response including contributions of synaptic depletion of neurotransmitters, strong hyperpolarizing currents in the soma preventing further spiking activity, small diameter (less myelination) of axons of cortical neurons, and GABA synaptic transmission [46–49]. The decline in activity at higher frequencies occurred in the model largely due to GABAergic synaptic transmission, and the inhibition lasted for several hundred milliseconds post-stimulation.

### Study limitations

Our cortical column model comprising neurons with realistic morphology and synapses is the most advanced modeling work to date to study the temporal effects of ICMS but nonetheless has some important limitations including modeling only a single column, lack of horizontal connections between columns, no connectivity between neurons, missing neural types, and lack of glia or other support cells that may contribute to the response. Despite these limitations, we replicated the major components seen in the experimentally measured temporal response to single pulse ICMS. However, we did not observe the post-inhibitory rebound excitatory response in the model neurons typically seen in the experimental response at the higher stimulation intensities and frequencies. The thalamus, which was not included in our model, appears to play a prominent role in generating the rebound response [50, 51]. Stimulating the cortex causes brief excitation followed by inhibition in the cortical neurons [51]. This excitation-inhibition response propagates to the thalamus from the cortex [51]. Rebound excitation is generated in the thalamus in the post-inhibitory phase due to the activation of the T-type Ca2+ channels and subsequently propagates back to the cortex [51]. Thalamic ablation suppressed the rebound excitation response in the cortical neurons [50]. Thus, the absence of the thalamus and corticothalamic and thalamocortical connections may explain the lack of rebound excitation in model neurons.

## Acknowledgements

The authors would like to thank Joseph Sombeck and Aman Aberra for helpful discussions on this work.

## Author Contributions

K.K. and W.M.G. conception and design of research; K.K. implemented computational model and ran simulations; K.K. analyzed data; K.K., and W.M.G. interpreted results; K.K. prepared figures and drafted manuscript; K.K. and W.M.G. edited and revised manuscript; K.K. and W.M.G. approved final version of manuscript.

## Supplementary Figures

**Figure S1:**
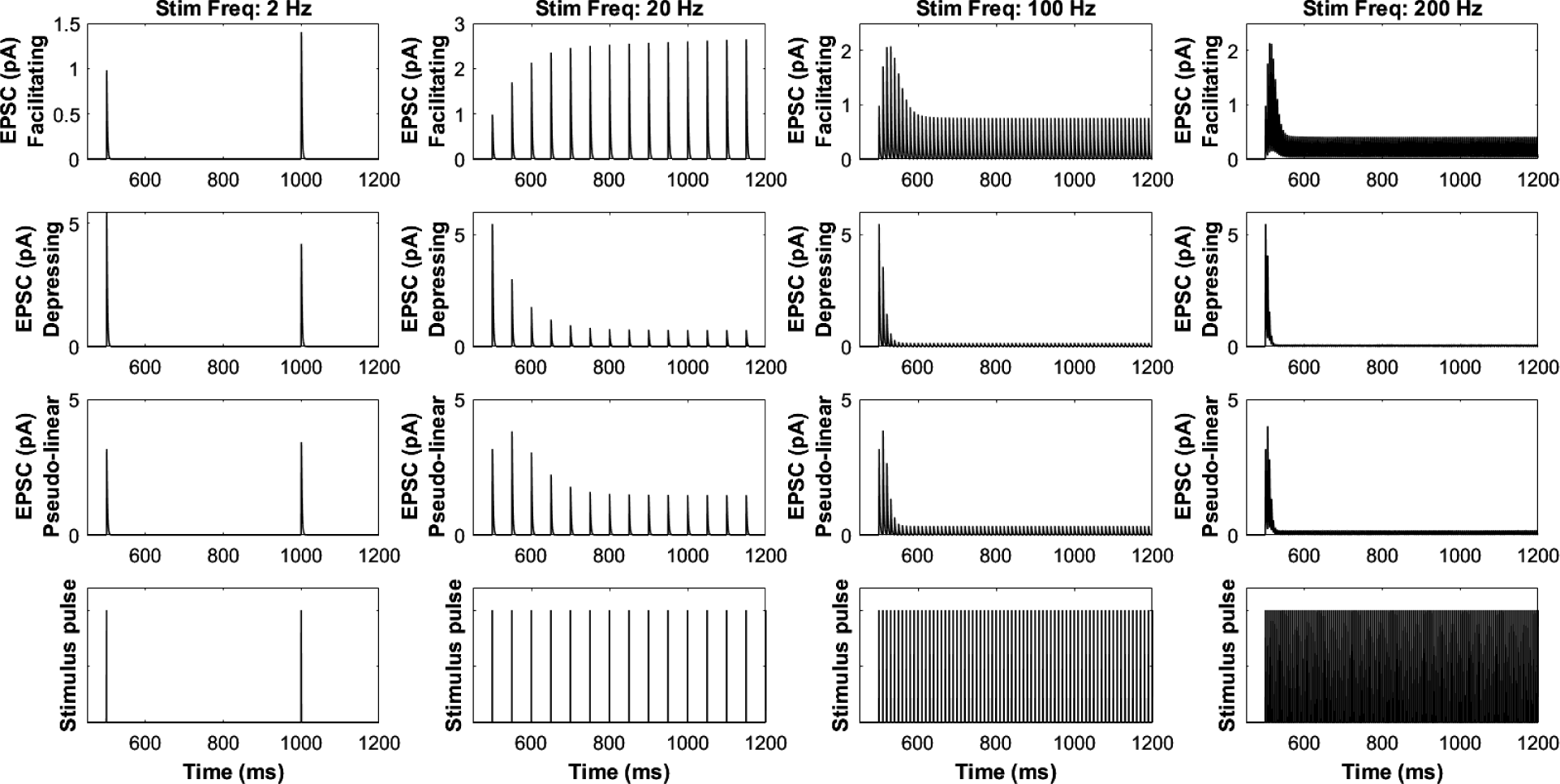
Properties of the (Tsodyks-Markram) AMPA synapse. Excitatory postsynaptic current (EPSC) showing dynamics of facilitating, depressing and pseudo-linear synapse in response to low (2, 20 Hz) and high (100, 200 Hz) frequency stimulation.

**Figure S2:**
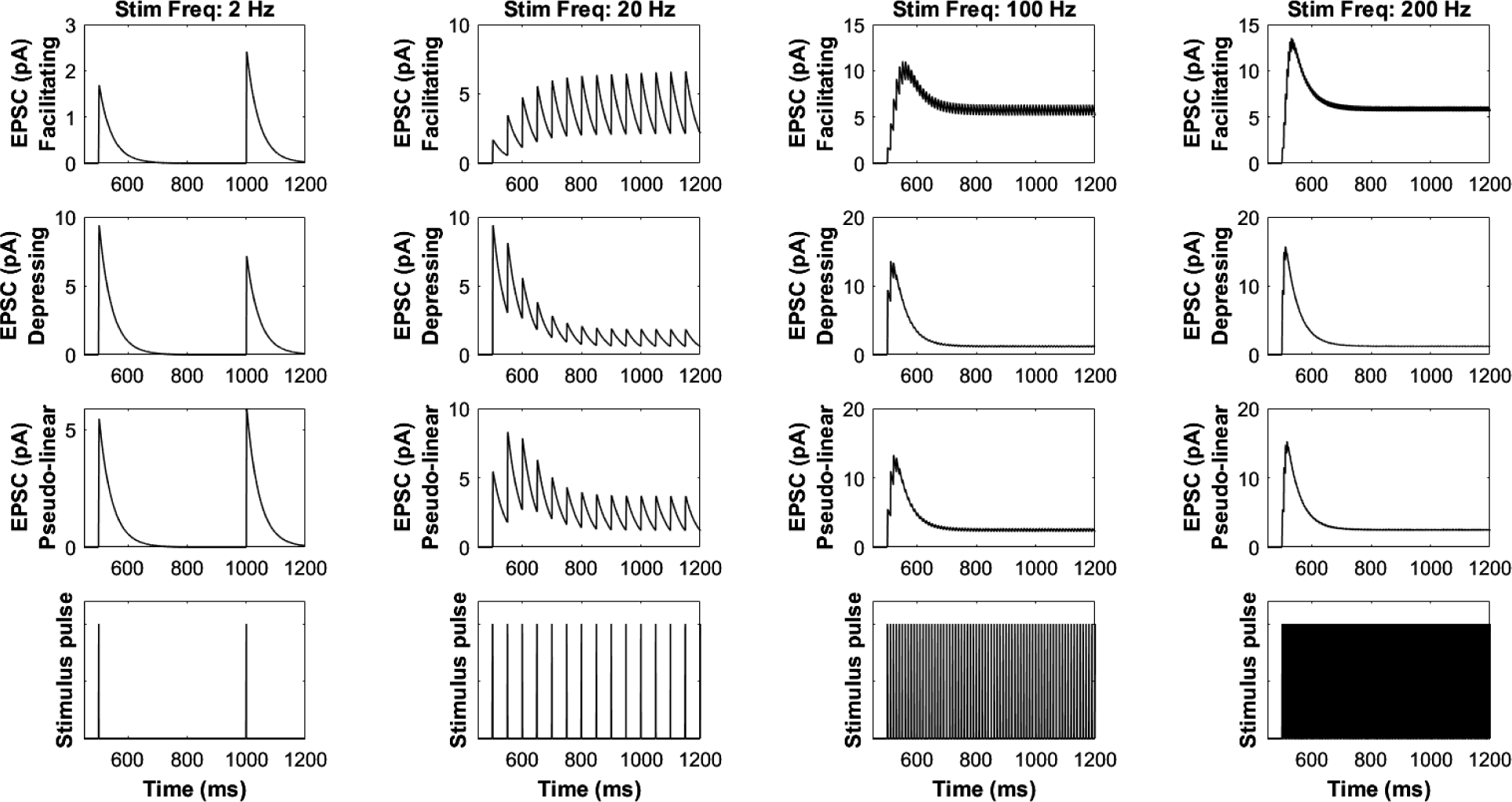
Properties of the (Tsodyks-Markram) NMDA synapse. Excitatory postsynaptic current (EPSC) showing dynamics of facilitating, depressing and pseudo-linear synapse in response to low (2, 20 Hz) and high (100, 200 Hz) frequency stimulation.

**Figure S3:**
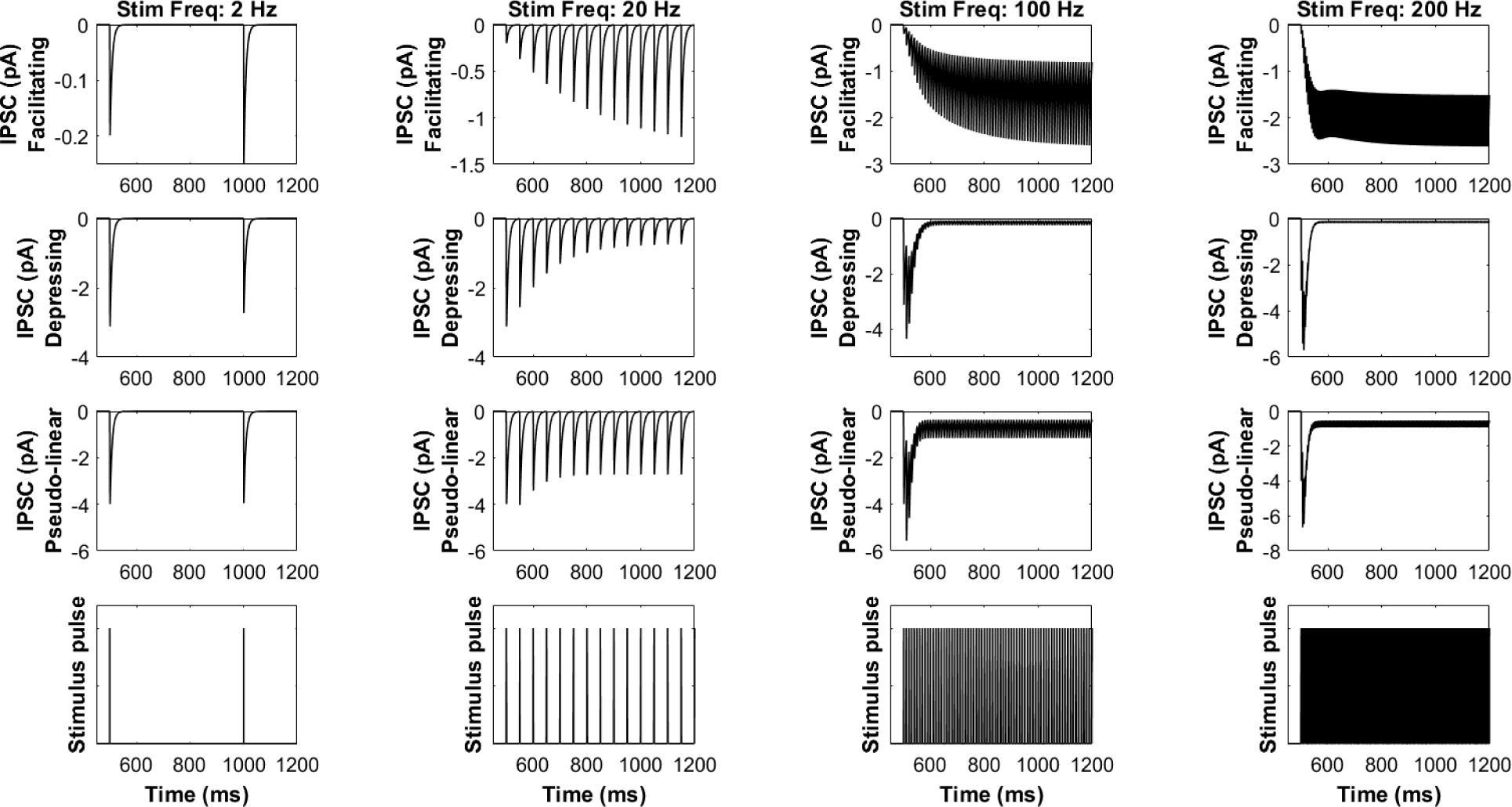
Properties of the (Tsodyks-Markram) GABA-A synapse. Inhibitory postsynaptic current (IPSC) showing dynamics of facilitating, depressing and pseudo-linear synapse in response to low (2, 20 Hz) and high (100, 200 Hz) frequency stimulation.

**Figure S4:**
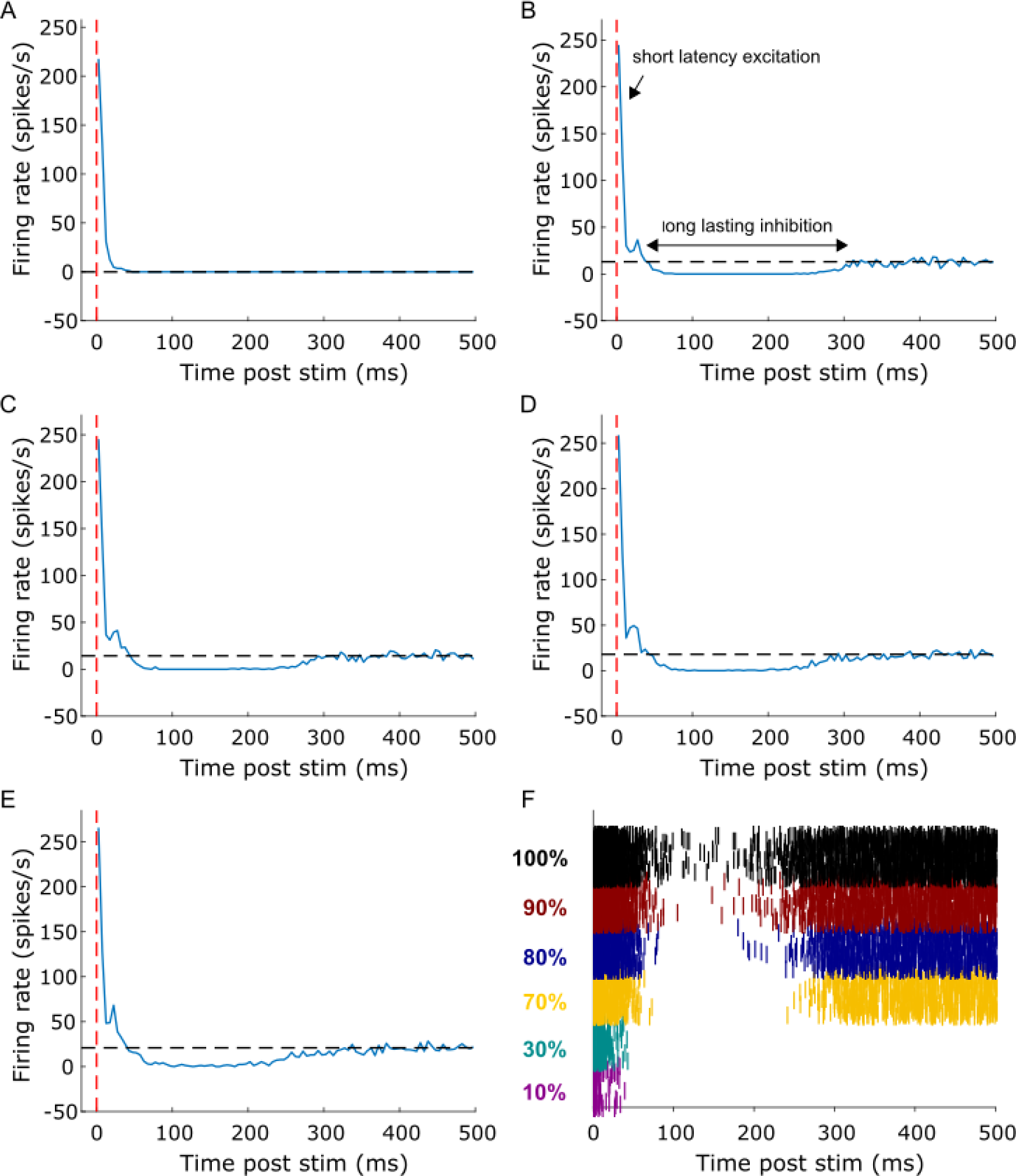
Effect of activating different L5 PC excitatory synapse proportions on the temporal response to ICMS. Temporal response to 50 µA ICMS at (A) 10%, (B) 70%, (C) 80%, (D) 90%, (E) 100% activation of excitatory synapses on L5 PC by 20 Hz Poisson spikes. Activation of 10% of excitatory synapses did not generate any intrinsic activity in the model neurons. The intrinsic firing rate increased marginally with a higher proportion of activation of synapses. (F) Raster plot showing firing activity of individual L5 PCs across different % activation of synapses.

**Figure S5:**
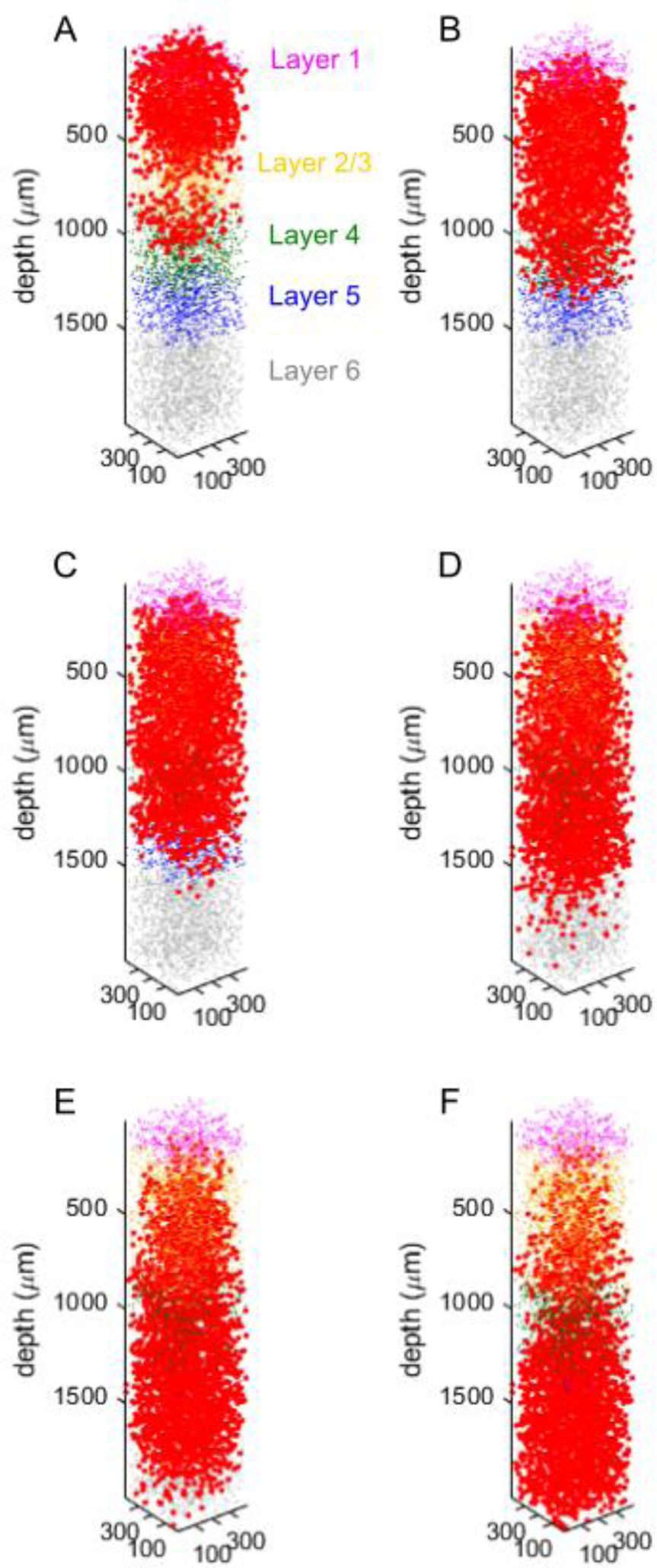
Response to ICMS to stimulation at different depths along the cortical column. (A) L1 depth: 300 μm, (B) L2/3 depth: 600 μm, (C) L4 depth: 900 μm, (D) L5 depth: 1200 μm, (E) L6 depth: 1500 μm and (F) L6 depth: 1800 μm. Each red dot represents the location of activated soma to ICMS. Stimulation comprised a single biphasic pulse (cathodic first) with a fixed width of 200 µs/phase, interphase interval of 50 µs and intensity of 30 µA. The 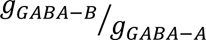 ratio was set at 1.

**Figure S6:**
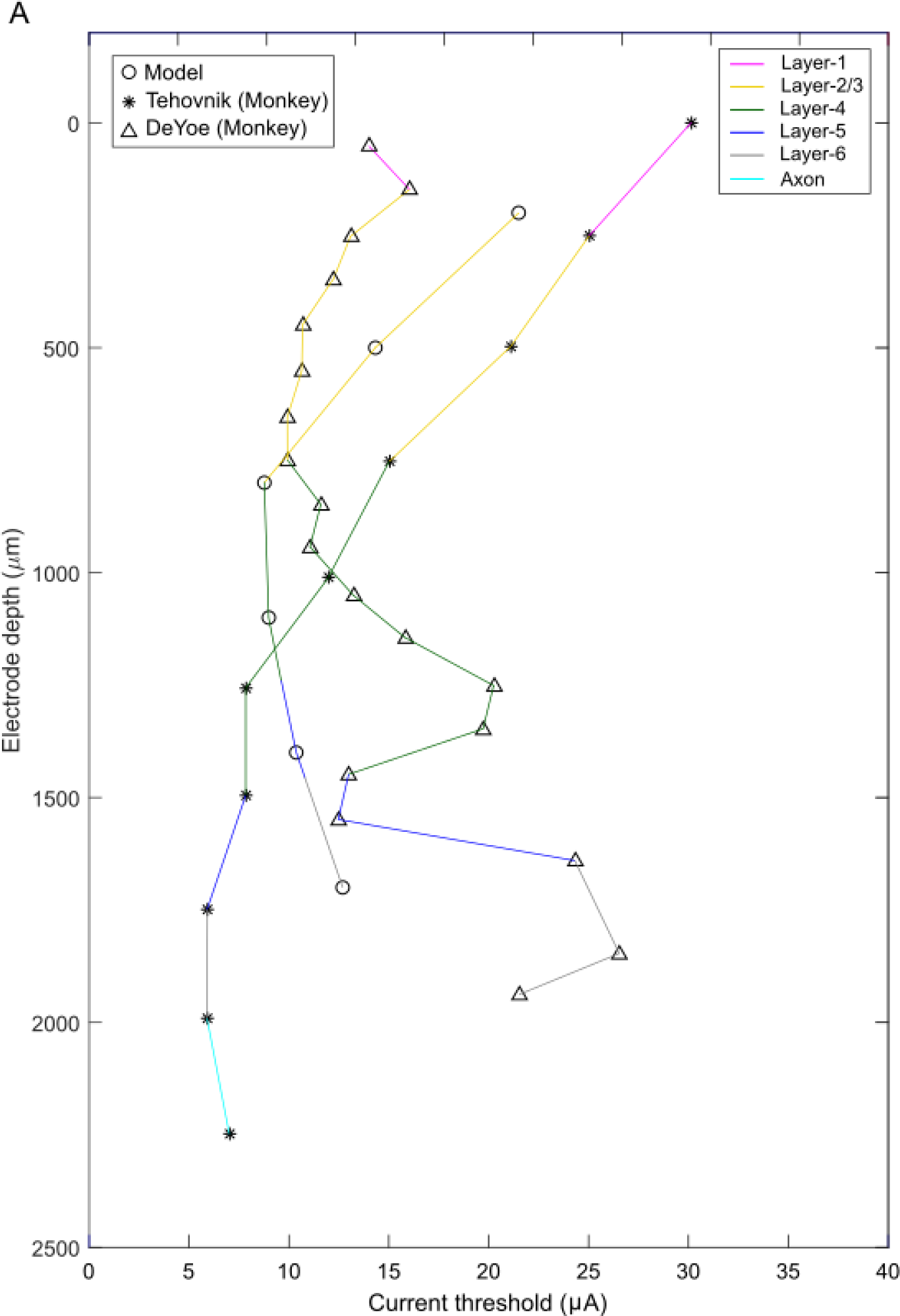
Depth**-**dependent effects of ICMS intensity. (A) Comparison of model-based current thresholds with detection thresholds from two non-human primate studies from the literature [55, 56]. The model’s current threshold at each depth was the lowest intensity that activated 1000 neurons. The current threshold was lower during stimulation in deeper layers compared to superficial layers.

**Figure S7:**
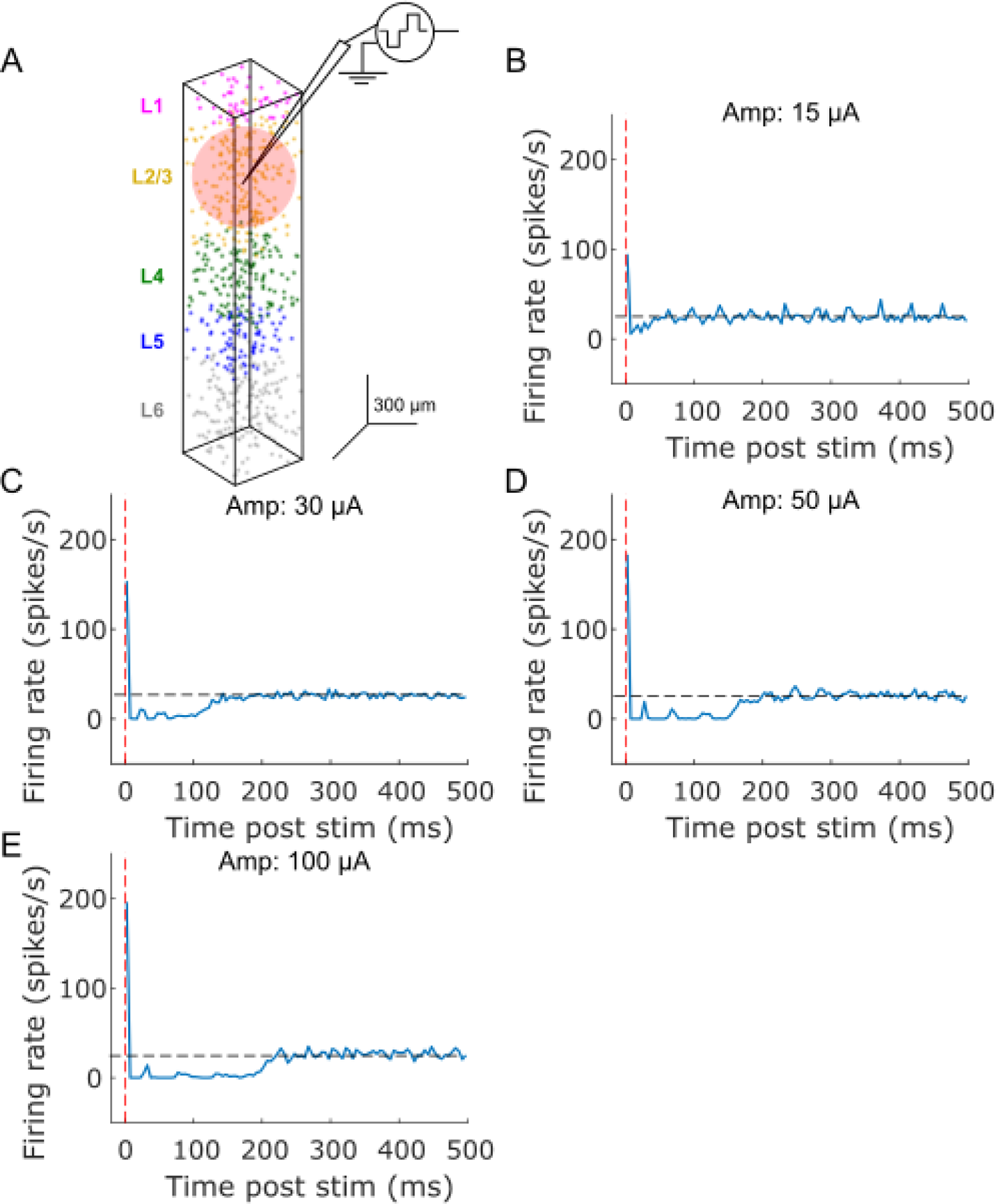
Temporal response of L2/3 pyramidal neurons. (A) Stimulating electrode was in L2/3. Temporal response of model neurons to ICMS at (B) 15 µA, (C) 30 µA, (D) 50 µA, (E) 100 µA. The response was averaged across L2/3 PC with somas located within 150 µm from the stimulation electrode (pink sphere in A). A bin width of 5 ms was used to bin spike times.

**Figure S8:**
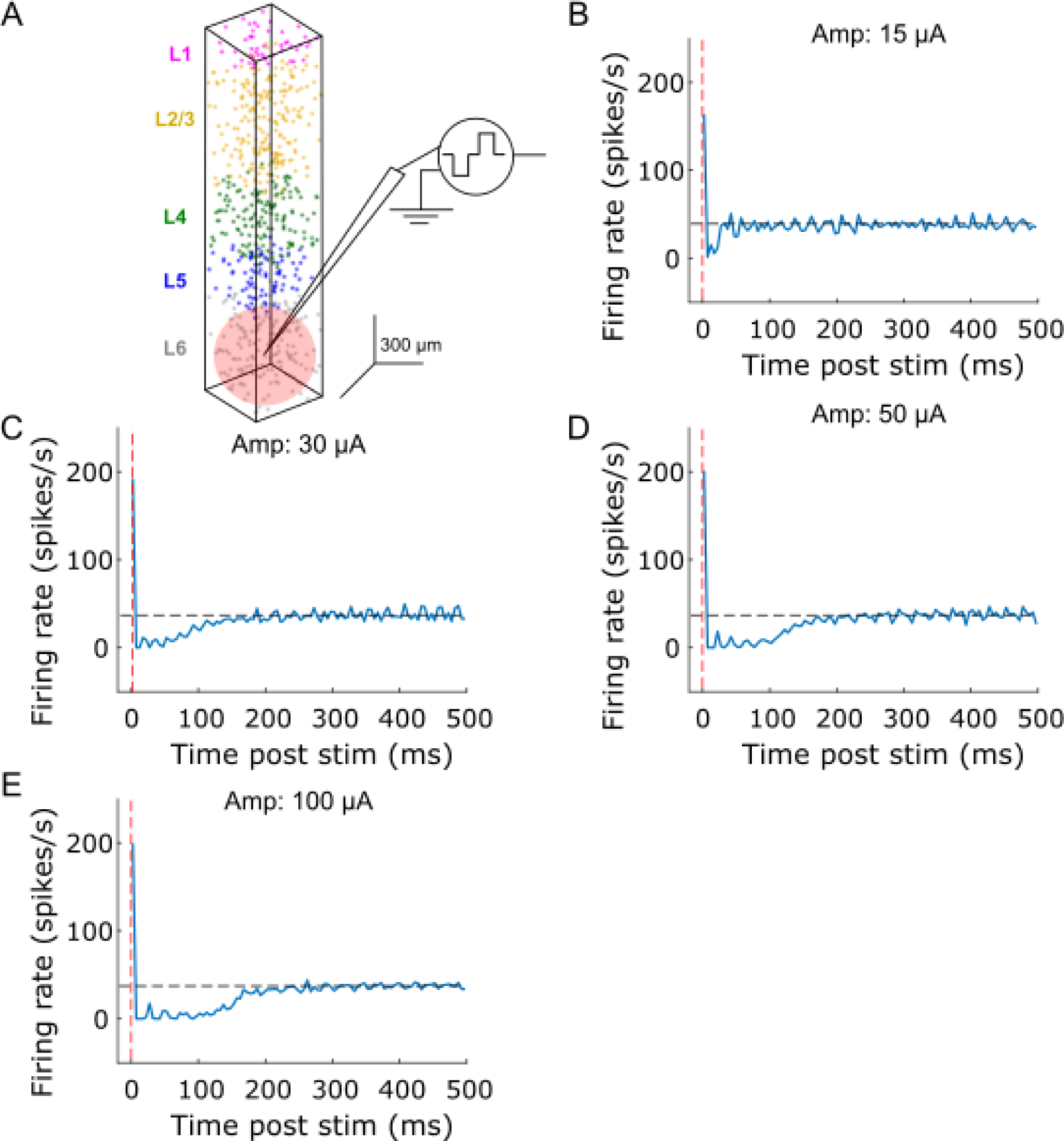
Temporal response of L6 pyramidal neurons. (A) Stimulating electrode was in L6. Temporal response of model neurons to ICMS at (B) 15 µA, (C) 30 µA, (D) 50 µA, (E) 100 µA. The response was averaged across L6 PC with somas located within 150 µm from the stimulation electrode (pink sphere in A). A bin width of 5 ms was used to bin spike times.

**Figure S9:**
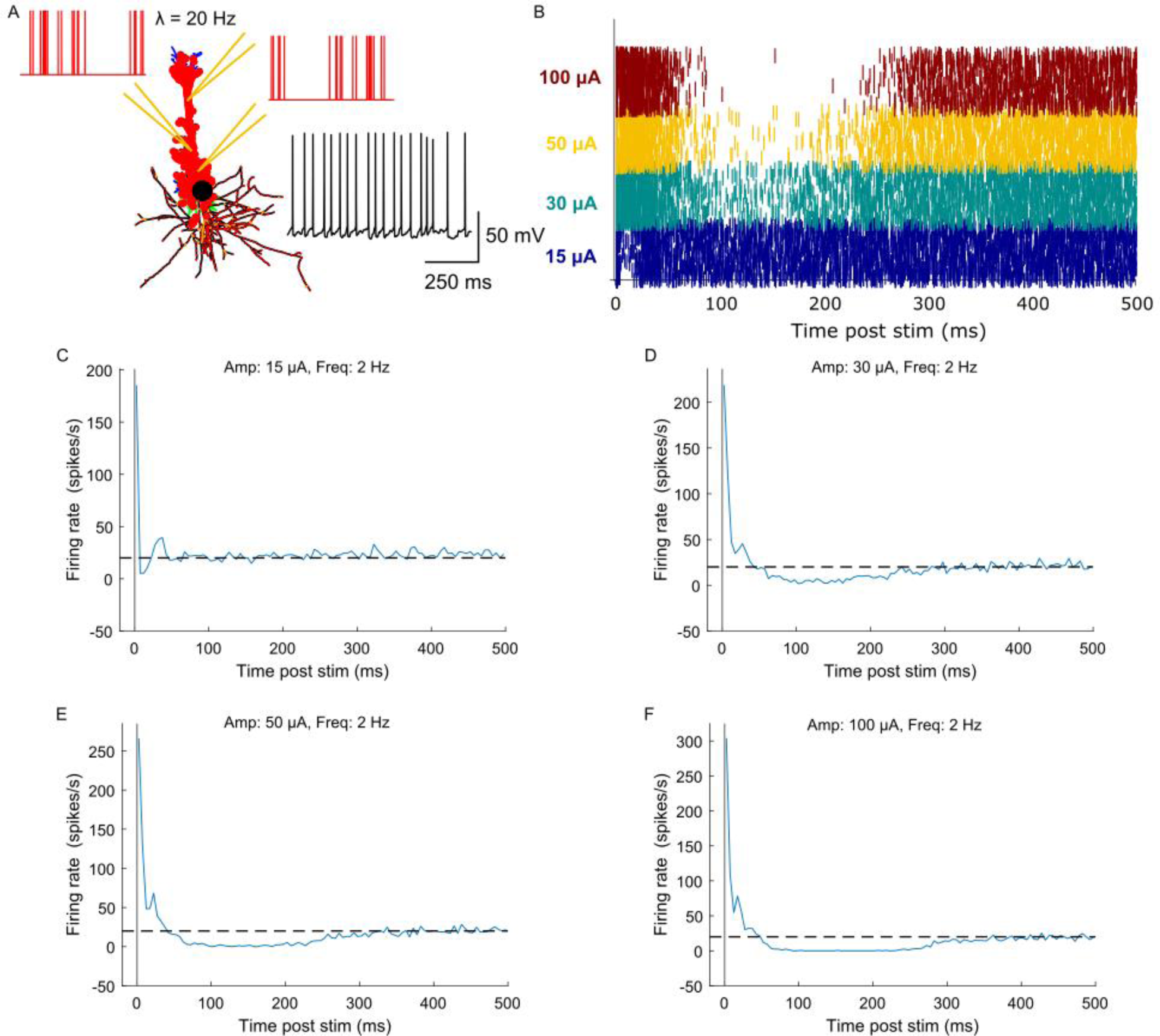
Temporal response of model neurons to intracortical microstimulation (ICMS) at different stimulation intensities and with intrinsic activity generated by activating excitatory synapses with Poisson spike trains. (A) The intrinsic activity (black trace) was generated by driving 90% of excitatory synapses on L5 PC (red-filled circles) with Poisson spikes at a mean rate of 20 Hz. (B) Raster plot showing firing activity of each L5 PC across stimulation intensities. Stimulation pulse triggered response to ICMS at (C) 15 µA, (D) 30 µA, (E) 50 µA and (F) 100 µA. A bin width of 5 ms was used to bin spike times.

**Figure S10:**
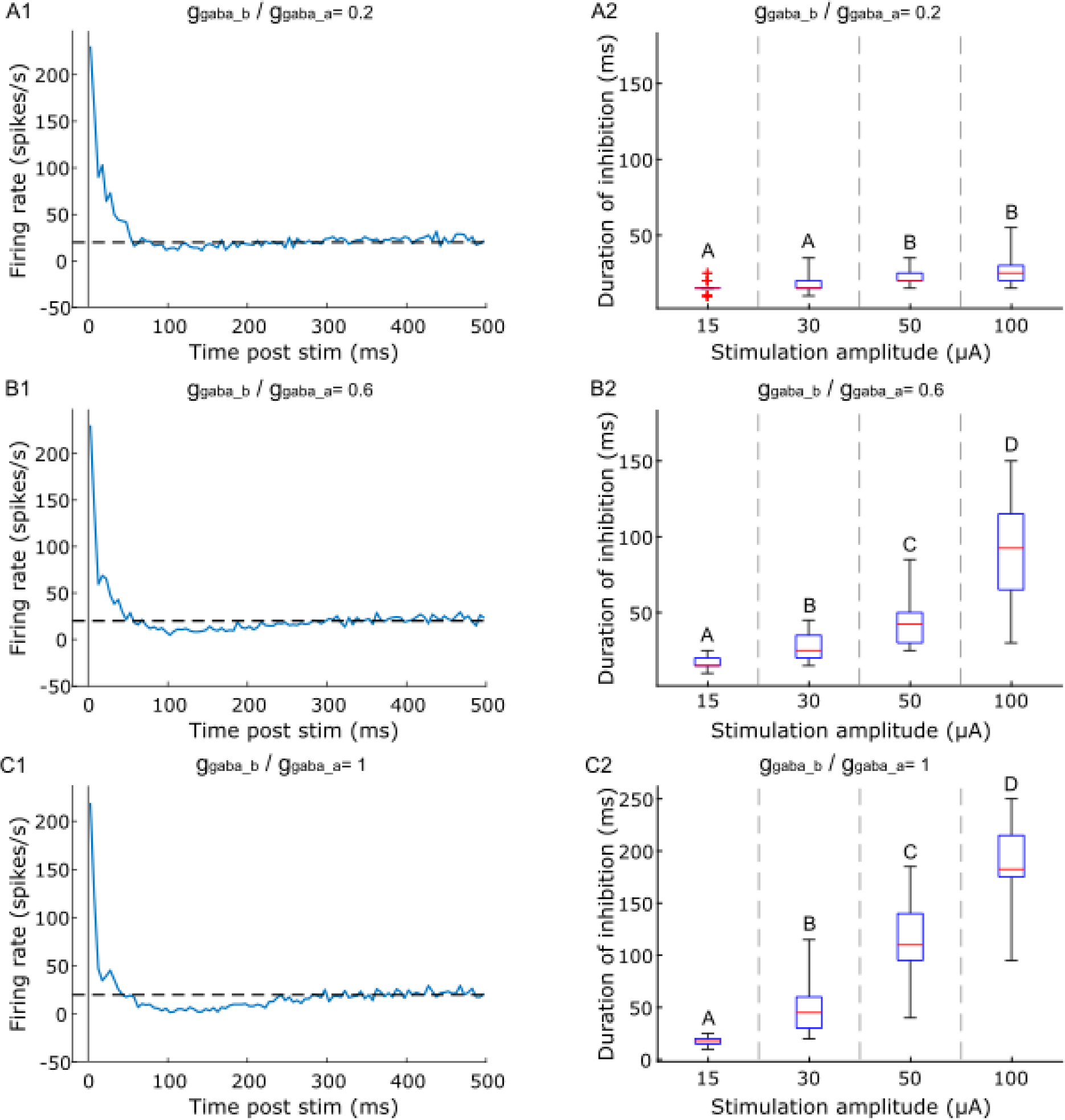
Response of model L5 PCs to L5 ICMS at 30 µA and 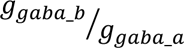 ratio of (A1) 0.2, (B1) 0.6, (C1) 1. The response was averaged across 34 L5 PCs with somas located within 150 µm of the stimulation electrode. A bin width of 5 ms was used to bin spike times. The intrinsic activity was generated by driving 90% of excitatory synapses on L5 PC with 20 Hz Poisson spikes. The duration of long-lasting inhibitory response as a function of ICMS amplitude for 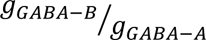 ratio of (A2) 0.2, (B2) 0.6, (C2) 1. Stimulation intensities that do not share the same letter are significantly different (p<0.05, Dunn-Sidak method). For each box, the central mark indicates the median duration across neurons, the bottom and top edges of the box indicate the 25th and 75th percentiles, the whiskers extend to 1.5 times the interquartile range and the plus signs indicate outliers.

**Figure S11:**
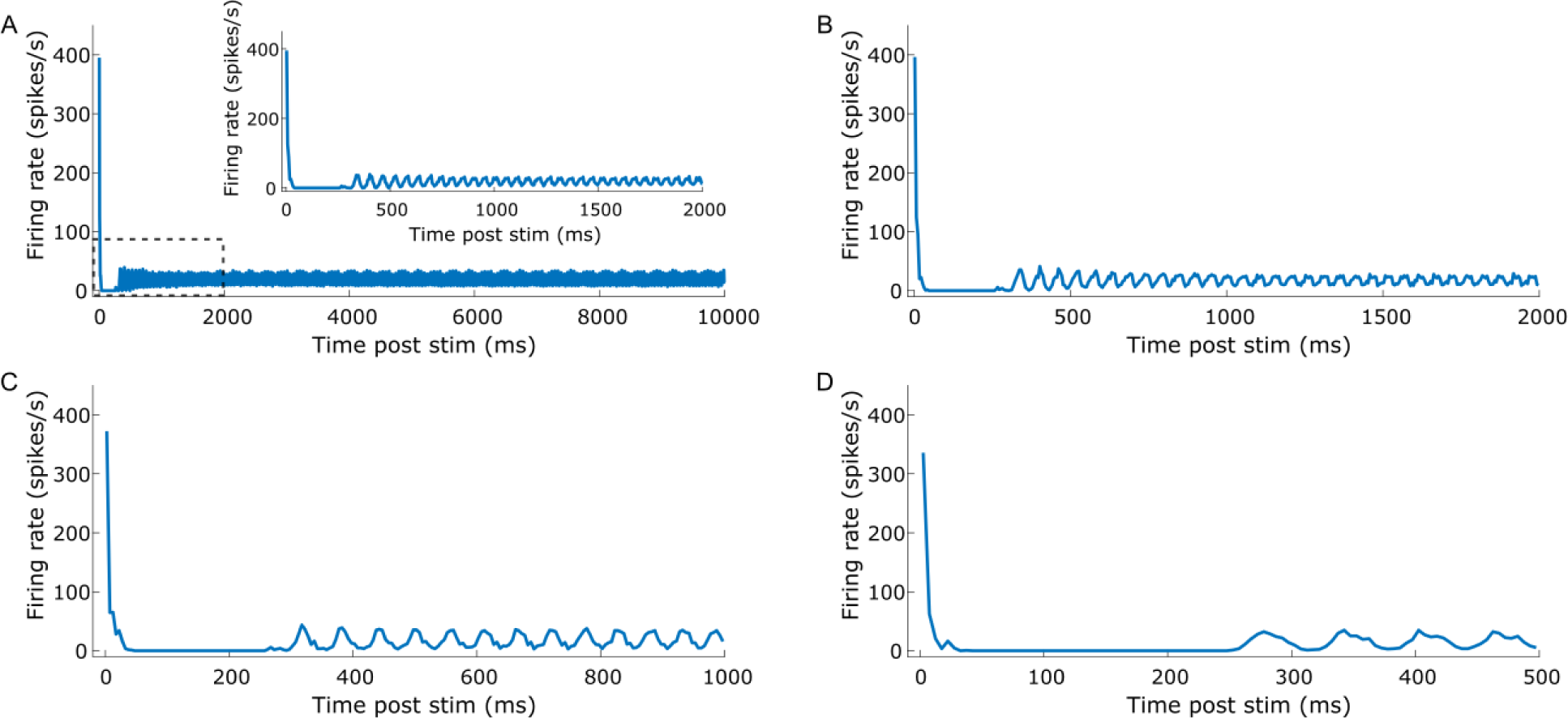
Stimulus triggered response to 100 µA ICMS pulse delivered at (A) 0.1 Hz, (B) 0.5 Hz, (C) 1 Hz and (D) 2 Hz. The inset in panel A shows the response to 0.1 Hz stimulation at a shorter time scale. For each condition, the stimulus triggered response was averaged across 10 pulses. The dashed line indicates the response without ICMS. The 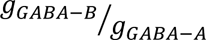 ratio was set at 1.

**Figure S12:**
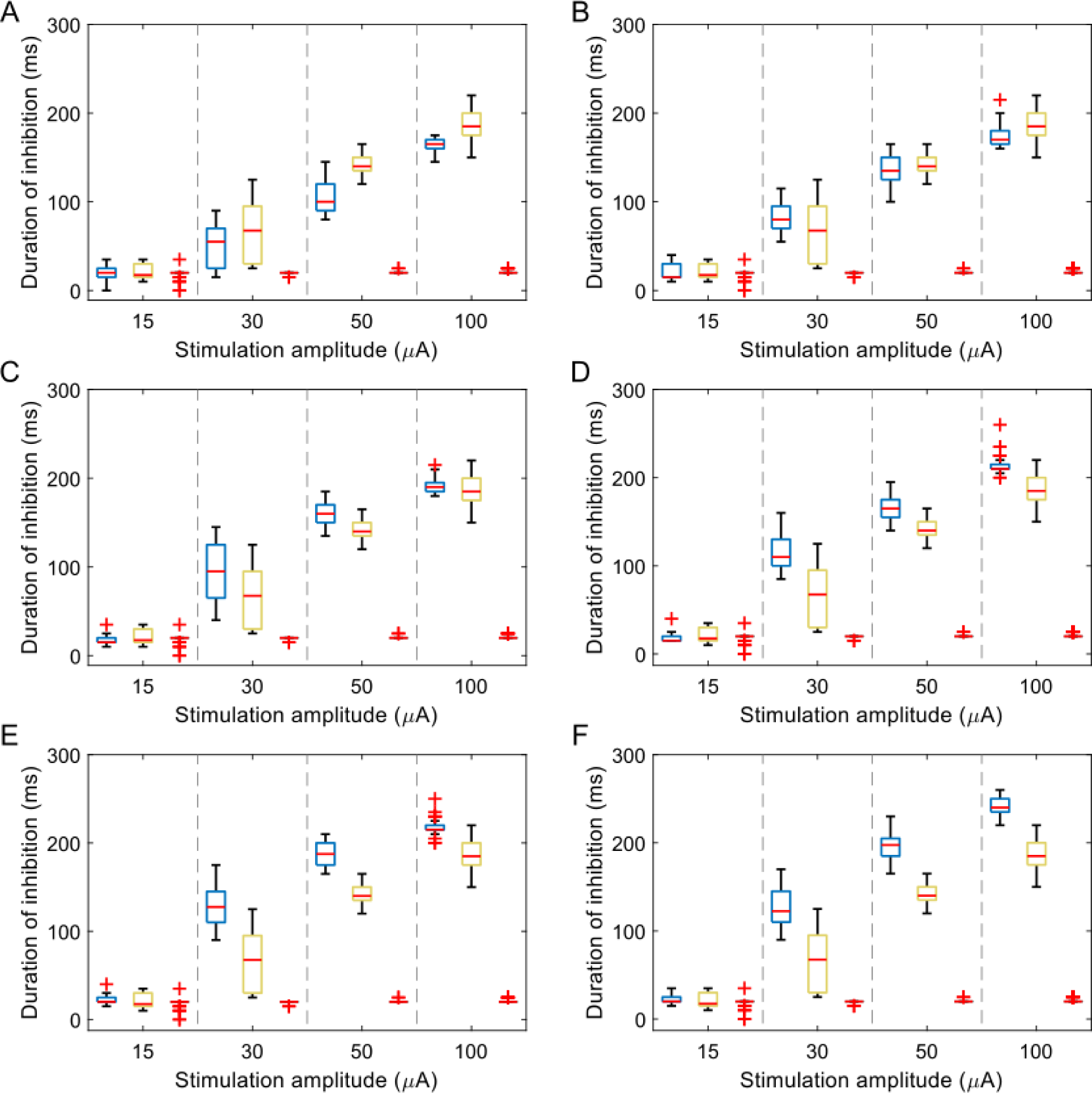
Duration of inhibition of model L5 PCs in response to L5 ICMS as a function of stimulation amplitude for 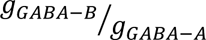 ratio of (A) 0, (B) 0.2, (C) 0.4, (D) 0.6, (E) 0.8, (F) 1. Each color box represents a different simulation condition: control condition (blue), without inhibitory synapses (yellow), without inhibitory and excitatory synapses (gray). For each box, the central mark indicates the median slope across neurons, the bottom and top edges of the box indicate the 25th and 75th percentiles, whiskers extend to 1.5 times the interquartile range and the plus signs indicate outliers.

**Figure S13:**
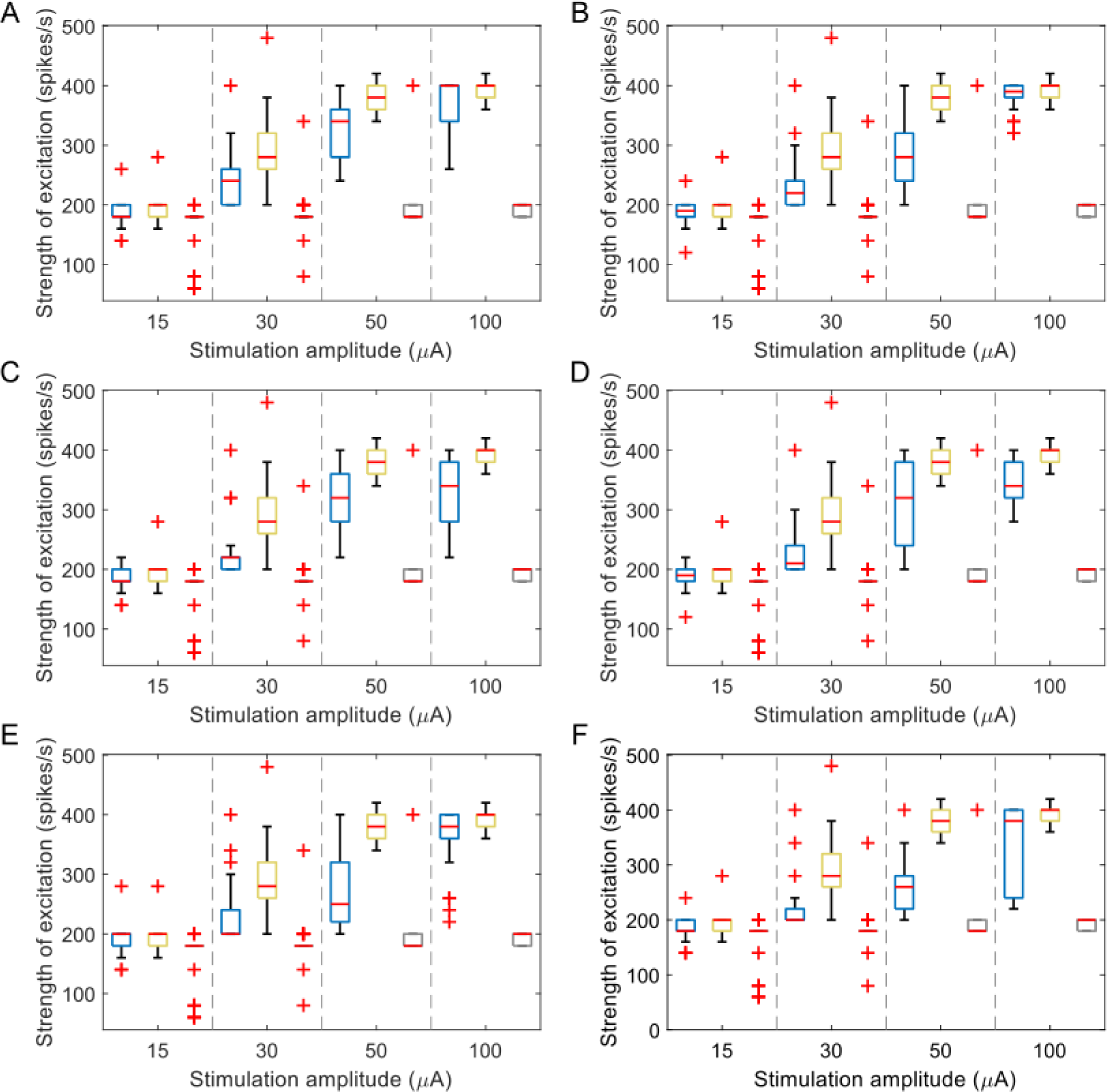
The peak of the excitatory response of model L5 PCs in response to L5 ICMS as a function of stimulation amplitude for 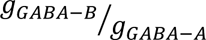 ratio of (A) 0, (B) 0.2, (C) 0.4, (D) 0.6, (E) 0.8, (F) 1. Each color box represents a different simulation condition: control condition (blue), without inhibitory synapses (yellow), without inhibitory and excitatory synapses (gray). For each box, the central mark indicates the median slope across neurons, the bottom and top edges of the box indicate the 25th and 75th percentiles, whiskers extend to 1.5 times the interquartile range and the plus signs indicate outliers.

**Figure S14:**
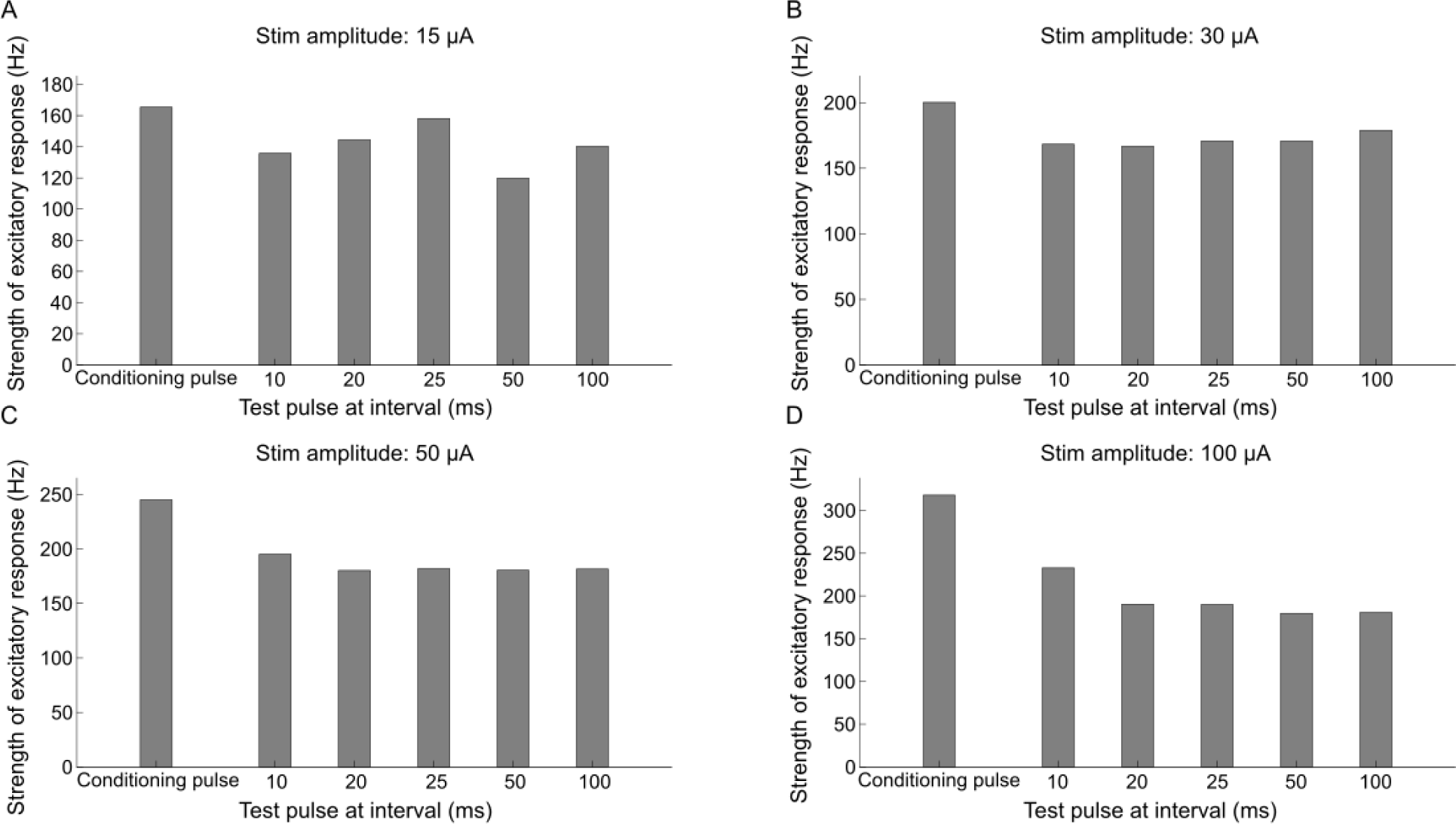
Influence of long-lasting inhibition on the excitatory response of model L5 PCs to L5ICMS. Strength of short latency excitatory response to test pulse at intensities: (A) 15 µA, (B) 30 µA, (C) 50 µA and (D) 100 µA. The conditioning pulse was applied at the same amplitude as the test pulse. The strength of excitatory response to test pulse decreased marginally compared to a single (conditioning) pulse and this effect was consistent across all stimulation intensities. This decrease can be attributed to the inhibitory effect triggered by the conditioning pulse.

**Figure S15:**
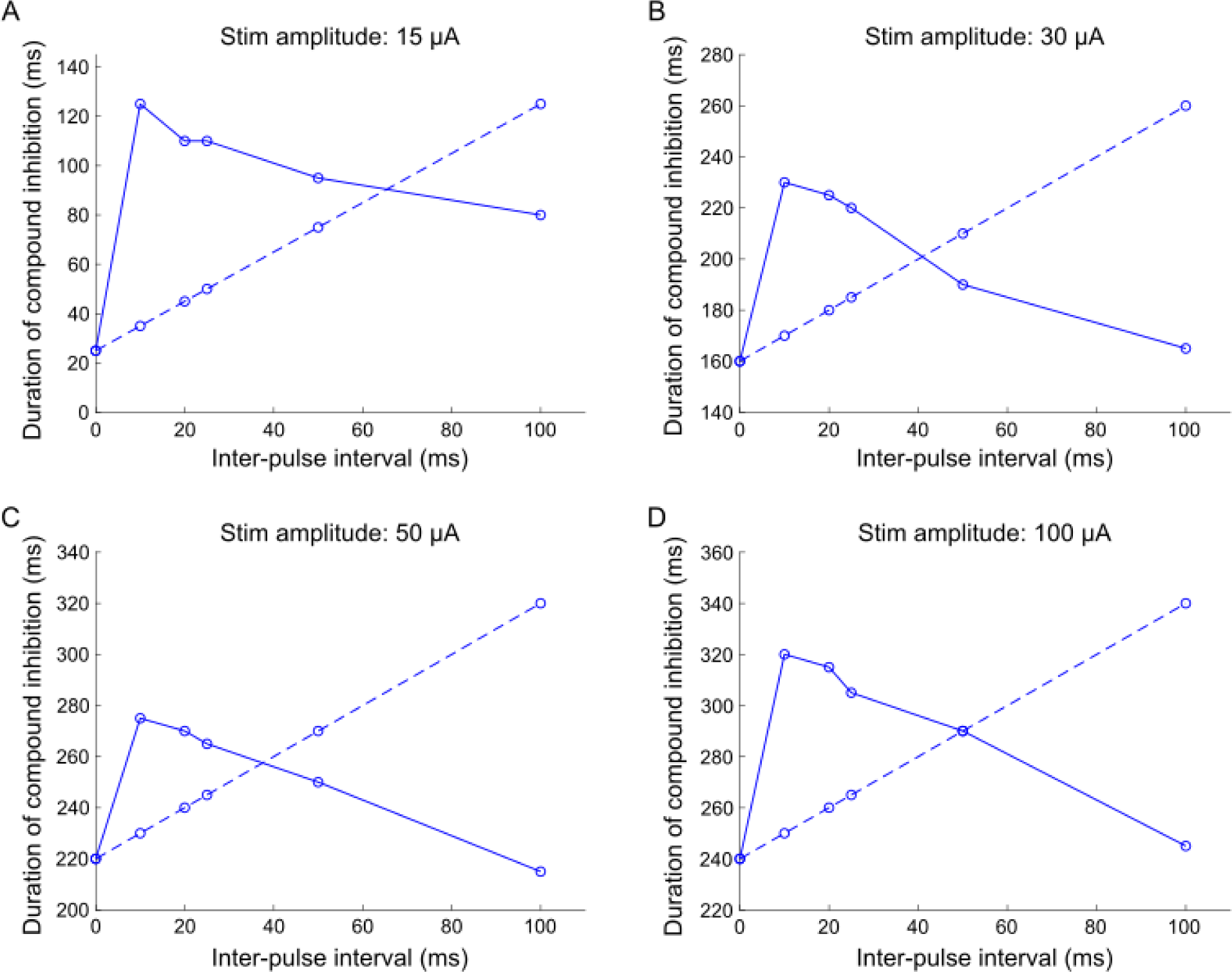
Interaction between inhibitory responses of model L5 PCs in response to during paired-pulse L5 ICMS. The duration of inhibitory response to test pulse at intensities: (A) 15 µA, (B) 30 µA, (C) 50 µA and (D) 100 µA. The conditioning pulse was applied at the same amplitude as the test pulse. Solid trace represents the inhibition duration to test pulse, whereas the dashed line indicates the duration predicted by linear superposition of individual inhibitory responses, i.e., duration to single pulse + IPI. There was supralinear addition of individual inhibitory responses for IPIs ≤ 40 ms and a sublinear superposition for IPIs >40 ms.

**Figure S16:**
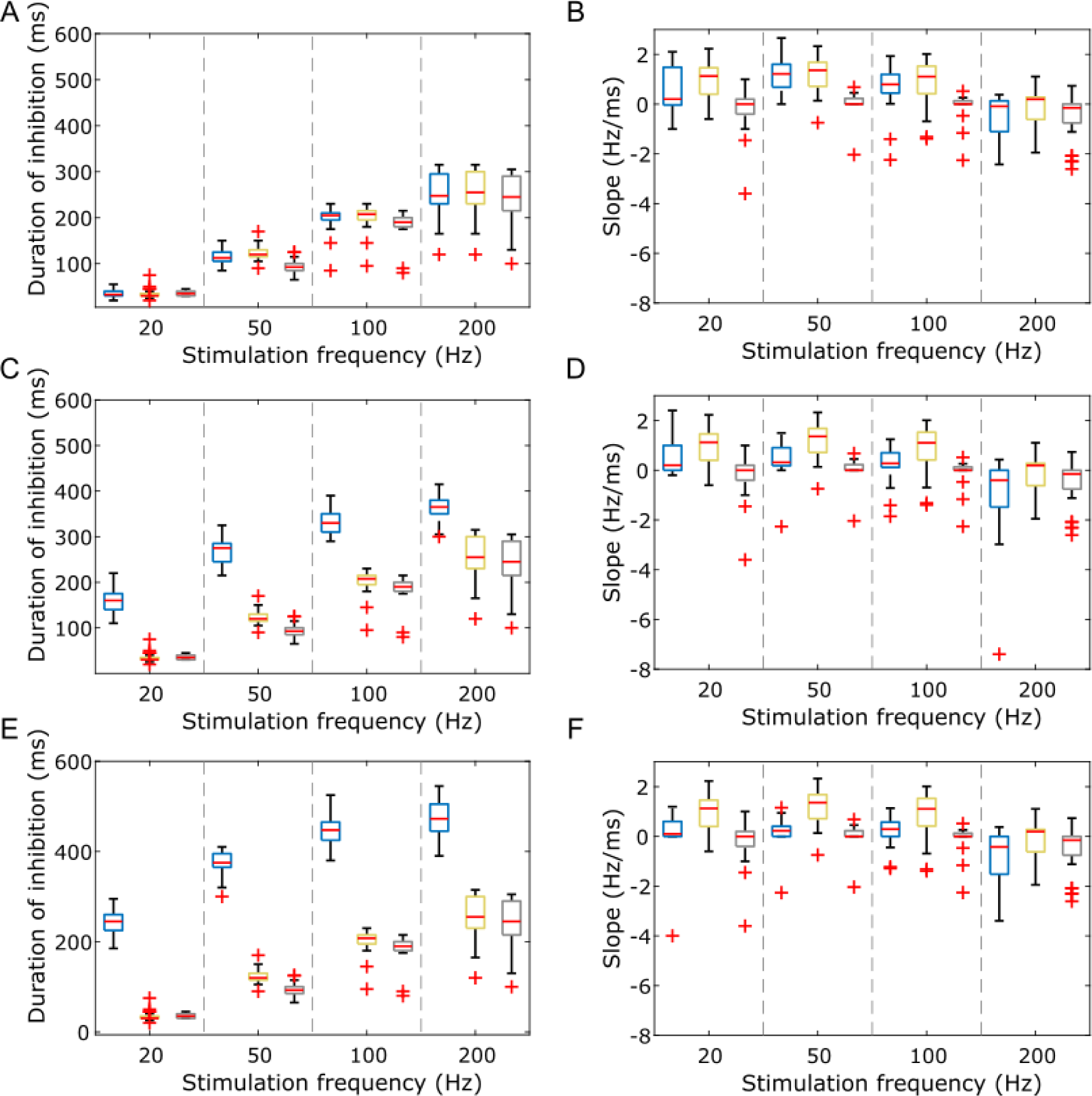
Duration of inhibition L5 PCs to L5 ICMS as a function of stimulation frequency for 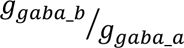 ratio of (A) 0, (C) 0.5, (E) 1. The slope of the line fit to the excitatory response as a function of stimulation frequency for 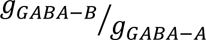 ratio of (B) 0, (D) 0.5, (F) 1. Each color box represents a different simulation condition: control condition (blue), without inhibitory synapses (yellow), without inhibitory and excitatory synapses (gray). For each box, the central mark indicates the median slope across neurons, the bottom and top edges of the box indicate the 25th and 75th percentiles, whiskers extend to 1.5 times the interquartile range, and the plus signs indicate outliers.

